# Neuromodulation in neural organoids with shell MEAs

**DOI:** 10.1101/2025.02.18.637712

**Authors:** Chris Acha, Derosh George, Lauren C. Diaz, Ziwei Ouyang, Dowlette-Mary M. Alam El Din, Hrishikesh Surlekar, Babak Moghadas, Eva Loftus, Gandhali M. Mangalvedhekar, Pratyush Sai R. Rayasam, Yu-Chiao Lai, Lena Smirnova, Brian S. Caffo, Erik C. Johnson, David H. Gracias

## Abstract

Neural organoids (NOs) have emerged as important tissue engineering models for brain sciences and biocomputing. Establishing reliable relationships between stimulation and recording traces of electrical activity is essential to monitor the functionality of NOs, especially as it relates to realizing biocomputing paradigms such as reinforcement learning or stimulus discrimination. While researchers have demonstrated neuromodulation in NOs, they have primarily used 2D microelectrode arrays (MEAs) with limited access to the entire 3D contour of the NOs. Here, we report neuromodulation using tiny mimics of macroscale EEG caps or shell MEAs. Specifically, we observe that stimulating current within a specific range (20 to 30 µA) induced a statistically significant increase in neuron firing rate when comparing the activity five seconds before and after stimulation. We observed neuromodulatory behavior using both three– and 16-electrode shells and could generate 3D spatiotemporal maps of neuromodulatory activity around the surface of the NO. Our studies demonstrate a methodology for investigating 3D spatiotemporal neuromodulation in organoids of broad relevance to biomedical engineering and biocomputing.

**One-Sentence Summary:** Neuromodulation, an essential intelligence feature, was observed using 3D stimulation and recording from neural organoids.

## INTRODUCTION

Recent studies have introduced three-dimensional (3D) self-aggregates of neural cells derived from human induced pluripotent stem cells (hiPSCs), known as neural organoids (NOs), as *in vitro* models of neural development, neurological disorders, and biocomputation (*1–8*). NOs have multiple cell types, including neurons, astrocytes, and oligodendrocytes with myelinated axons (*9*). Also, due to their 3D geometry and cellular interconnectivity, NO models can better recapitulate cellular heterogeneity and complexity than 2D culture models (*4*, *5*, *10*).

NOs have been widely studied using conventional microelectrode arrays (MEAs), including functional characterization, learning, neuronal circuitry, and functionality (*2*, *11–13*). However, a key limitation of conventional MEAs is that they are inherently 2D, limiting stimulation and recording to cells only on the bottom of the organoid. There are a few demonstrations of 3D MEA systems, which are wrapping NOs with self-folding or buckling sheets containing electrode arrays (*14*, *15*), penetrating needles-like electrodes into the organoids (*16*), and embedding electrode arrays in the culture during organogenesis (*17*). Many of these platforms are in early development, and neuromodulation using both stimulation and recording electrical signals has yet to be systematically realized.

We previously reported a self-folding MEA shell for recording the electrical activity of NOs (*14*). Here, we incorporate the capacity for both stimulation and recording to realize a platform for 3D neuromodulation from live NOs (**Fig. 1**). To accomplish this feature, we developed an integrated package so that continuous recording could be achieved while the NOs were inside an incubator. The MEA shell is modeled on *in vivo* recording systems such as electrocorticography (ECoG) arrays that enable surface recordings from the 3D surface of the brain (*18–20*). ECoG arrays characterize population dynamics, functional connectivity, and individual neuronal responses *in vivo*. They allow spatiotemporal mapping of population responses, characterization of high-gamma activity (*21*), and designing high-performing brain-computer interfaces (*22*).

**Fig. 1.**
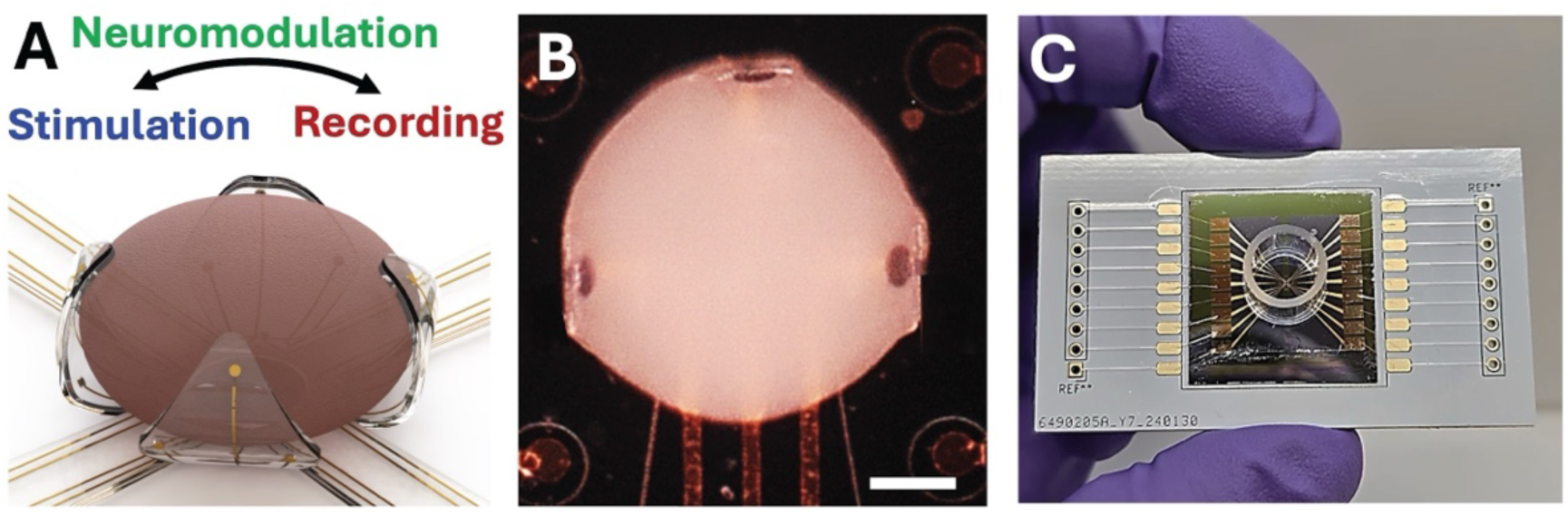
Conceptual schematic and integrated shell MEA used for 3D neuromodulation. **(A)** Schematic showing conformal wrapping of a MEA shell around the contour of a neural organoid. **(B)** Bright-field optical micrograph of a live NO within a self-folded MEA shell with three leaflets. Scale bar, 200 µm. **(C)** Image of a packaged shell NO device featuring the self-folding MEAs, cloning cylinder for culture media, bond pads, on and off-chip wiring, and packaging on a PCB board.

In this study, we used MEA shells to both stimulate and record electrical signals from around the 3D contour of live NOs using three-or 16-electrodes distributed around the spherical contour of the shell. Electrodes were distributed symmetrically across three– and four-leaflet designs to maximize spatial mapping around the spherical-like contour of the NO. We coated the contact pads with conductive poly-(3,4-ethylenedioxythiophene): poly (styrene sulfonate) (PEDOT: PSS) to reduce impedance for enhanced signal-to-noise ratio (SNR) and to increase the charge injection capacity (CIC) to prevent tissue damage during stimulation. We verified the biocompatibility of the shell electrodes by long-term recording and stimulation of organoids encapsulated inside these 3D shell electrodes for up to six weeks.

By systematically varying the amplitude of single stimulation pulses and analyzing correlations to recording traces, we discovered that it was possible to modulate the firing rate of single-unit neuronal recording traces. We also observed activity modulation to local field potential (LFP) in response to stimulation. We characterized spatial and temporal responses, demonstrating the ability to record single units independently and monitor electrical activity over weeks in culture. Finally, we developed a mathematical framework for spatiotemporal signal mapping from NOs using shell MEAs. This approach will advance neuromodulation with 3D spatiotemporal control of broad relevance to developmental biology, biomedical engineering, and biocomputing.

## RESULTS

### Design and fabrication of shell MEAs

We investigated 3D electrical neuromodulation by encapsulating NOs within fully packaged shell MEAs equipped with a cloning cylinder for cell media, 3D electrical stimulation and recording interfaces, and bonding pads (**Fig. 1**). We fabricated self-folding polymeric shell MEAs using a previously published protocol on silicon wafers with some modifications (*14*). The shell MEAs comprise a bilayer of differentially photo-crosslinked SU8 epoxy with metal (gold, Au) wiring and electrode contact pads coated with a conductive polymer (PEDOT: PSS). We fabricated the shell MEAs on insulating oxide-grown silicon wafers using photolithography. We utilized seven computer-aided design (CAD) masks (**Fig. S1-S3**) to define, (i) a sacrificial germanium (Ge) layer using the lift-off process, (ii) the first SU8 layer (exposed at 240 mJ/cm^2^ for three-electrode shell and 160 mJ/cm^2^ for 16-electrode shell MEA), which forms the base of the leaflets of the self-folding shell, (iii) Au wiring and electrode contact pads using a lift-off process, (iv) copper (Cu) bond pads around the shell for external interconnects and packaging using a lift-off process, (v) a second SU8 layer (exposed at 240 mJ/cm^2^ for three-electrode shell and 150 mJ/cm^2^ for 16-electrode shell MEA) to insulate the electrode wiring, (vi) a third SU8 layer (exposed at 120 mJ/cm^2^) which forms the top layer of the leaflets for bilayer self-folding, and (vii) selective SU8 regions (additionally exposed at 120 mJ/cm^2^) to tune the bending of the leaflets (**Fig. S1-S4**).

A highlight of our approach is that the fabrication process is chip-based, very large-scale integration (VLSI) compatible, and parallel, such that we could fabricate four shell MEAs simultaneously on a 3-inch wafer (**Fig. 2**), which is equivalent to about 64 shell MEAs being patterned on 12-inch diameter wafers used in commercial VLSI chip foundries. It is also possible to customize shapes and electrode configurations (**Fig. S5**). After fabrication, we diced the wafer into individual chips, each consisting of a single MEA shell and peripheral bond pads. We then attached the silicon chip to an in-house designed printed circuit board (PCB) using polydimethylsiloxane (PDMS) and wire-bonded the peripheral bond pads to the PCB carrier (**Fig. S6**). Finally, we attached a cloning cylinder around the shell and sealed it using PDMS.

**Fig. 2.**
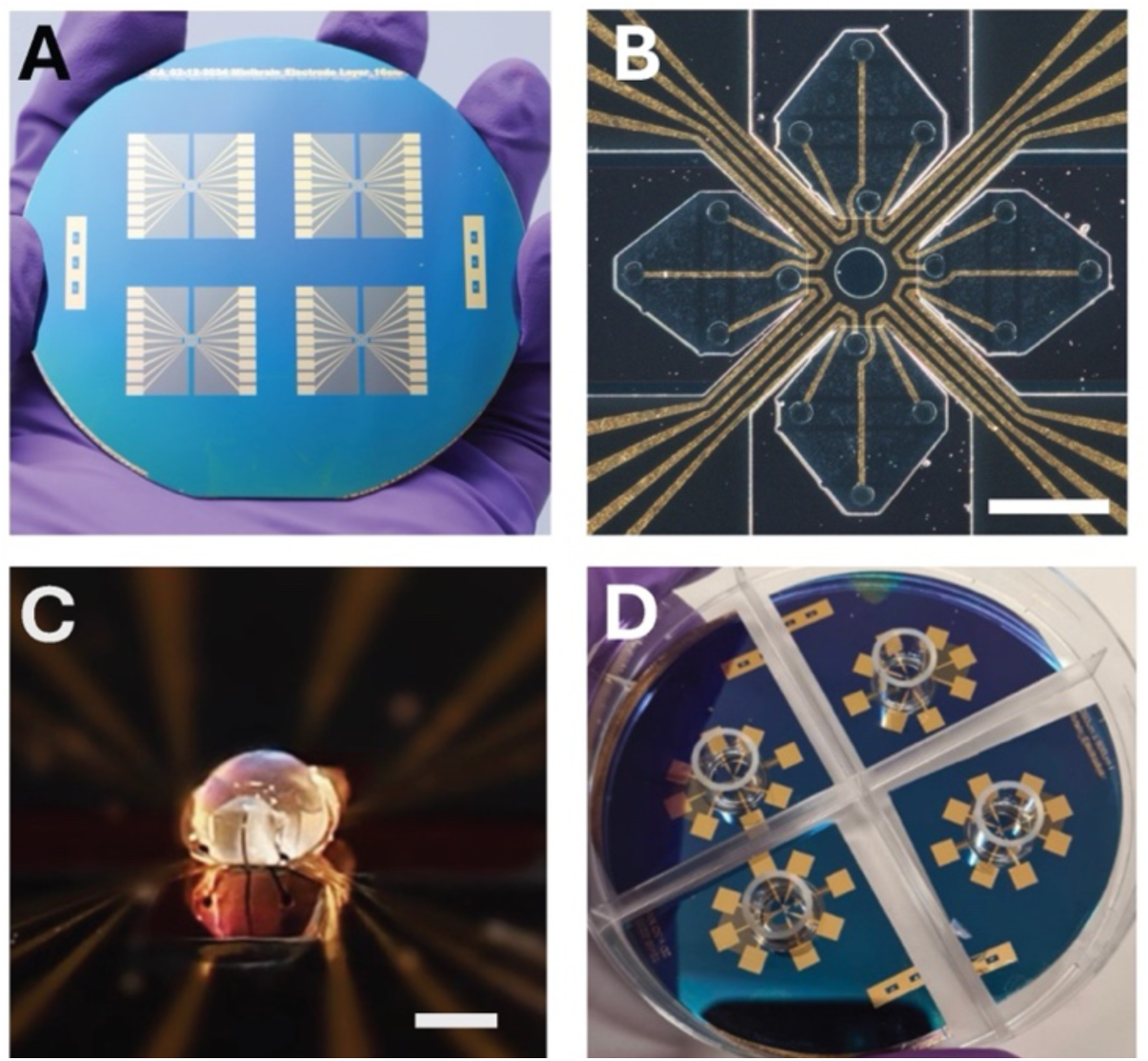
Fabrication and self-folding process. **(A)** Image of shell MEA and wiring patterned using seven-layer CAD-based photolithography. **(B)** Zoom-in micrograph of a single pre-folded shell MEA with four leaflets, 16 electrodes with gold (Au) wiring and conductive polymer pads on a silicon wafer. Scale bar, 500 µm. **(C)** Image of a self-folded shell MEA encapsulation of an alginate bead. Scale bar, 500 µm. Self-folding was triggered on placement in water. **(D)** Image of four shell MEAs with packaged glass cylinders to hold media illustrating that the fabrication process enables customization and mass-production.

### Electrochemical characterization of PEDOT: PSS contact pads in the shell MEAs

We electrodeposited conductive PEDOT: PSS onto the shell electrode contact pads to decrease electrode impedance, improving the signal-to-noise ratio in neural recordings (*23*). We observed using electrochemical impedance measurements that a 10-μm thick film lowered the impedance of the 60-μm diameter gold contact pads by about 98% (from 222 kΩ to 4 kΩ) at 1 kHz with consistent performance (**Fig. S7**). We also performed cyclic voltammetry (CV) at a 200-mV s^−1^ scan rate to measure the contact pads’ charge storage capacity (CSC). We found an improved CSC of 69 mC/cm^2^ for PEDOT: PSS-coated contact pads compared to 0.36 mC/cm^2^ for bare Au pads (**Fig. S8**). This increase in CSC indicates that a PEDOT: PSS-coated contact pad can inject more charge during stimulation than bare Au pads (*23*). We also measured voltage transient response to stimulation pulses and showed that PEDOT: PSS-coated electrodes’ charge injection capacity (CIC) is higher than that of bare Au pads (**Fig. S9**). A higher charge storage and injection capacity enables the application of a broader range of stimulating currents without causing the electrolysis of water (**Fig. S9**) (*23*). Furthermore, PEDOT: PSS-coated Au electrodes had lower polarization potentials during stimulation than bare Au electrodes, indicating that PEDOT: PSS coating can increase the durability of the Au electrodes, like preventing corrosion or delamination caused by undesired electrolysis.

### NO characteristics

We generated NOs from hiPSCs and cultured them for 4–8 weeks using an unguided differentiation protocol. This protocol yields NOs of controlled size and shape, minimizing necrosis in their centers. Previous studies have shown that this model produces NOs with an average diameter of approximately 300-600 μm from weeks 3-12 after initiating neural maturation in 3D (*9*). NOs generated using this protocol consist of main cell types found in the central nervous system (*9*, *24*, *25*). **Fig. S10** shows the expression of mature and immature neuronal and dendritic markers. In previous studies, time-course characterization of NO differentiation has shown increasing expression of markers for oligodendrocytes, astrocytes, and neurons over time, accompanied by a decrease in nestin, a progenitor cell marker. Specifically, neuronal populations, including GABAergic, glutaminergic, and dopaminergic neurons, have been identified in this model (*9*, *24*, *25*).

Furthermore, previous studies have shown that such NOs can resemble specific human brain regions. RNA-seq data from week 8 NOs showed expression profiles resembling the temporal neocortex and ventrolateral prefrontal cortex (*24*, *26*). From time-course analyses of the electrical activity of NOs, we observed increasing spontaneous electrical activity over differentiation (*24*). Quantification of electrophysiology data revealed highly connected neuronal networks, with connectivity and criticality increasing over time, reflecting the maturation of these systems (*24*). Additionally, we observed evoked activity alongside increased expression of immediate early genes (IEGs) following electrical or pharmacological modulation of GABAergic and glutamatergic neurons with eliciting network spiking. These characteristics further confirm the functional maturation of these NOs (*14*, *24*).

### Long-term NO electrical recording characteristics using shell MEAs

We used the following protocol for long-term NO spatial recording. Briefly, we sorted NOs based on size and age of differentiation. We encapsulated live NOs within the self-folding shells in culture media in the cloning cylinder packaged on a printed circuit board (PCB). We developed and utilized a fully integrated workstation using the Intan RHS Stim/Recording System to enable continuous electrical interrogation within the incubator. We placed the NOs in the incubator during recording with a three-electrode (North, West, and East) shell layout distributed around a sphere. While the NOs were encapsulated within the shell MEAs, we changed the neural differentiation culture media every two to three days to maintain organoid health and ensure consistent data collection. We recorded multichannel electrophysiology data using a multichannel amplifier and data acquisition system. We conducted all recordings at a sampling rate of 30 kHz, allowing for fast-paced, high-resolution, long-term observations of any spiking that occurred, and measured both single-unit spikes and LFPs. To eliminate the natural AC outlet oscillation, we applied a 60 Hz notch filter and a high-pass filter to detect spikes at a default relative threshold of 5.5 times the root mean square (RMS) voltage (which was then adjusted offline to account for signal SNR during spike sorting, see **Note S1**).

We analyzed the data using an open-source Python package, SpikeInterface (*27*), including bandpass filtering of the signal for analysis of specific frequency bands and spike sorting of single units. We rejected stimulus artifacts using a threshold-based approach and performed spike sorting with the package MountainSort5 (*28*). Two researchers reviewed the results to remove noise units and resolve split clusters. Our electrophysiological analysis pipeline can sort detected spikes and evaluate LFPs. In the Methods section, we provide additional details and parameters on the spike sorting approach (**Note S1**). We first verified that the 3D MEA recording signals were qualitatively consistent with 2D MEA raw recordings and spike-time raster plots (**Fig. 3 and S11**), which provided evidence that signals recorded by the 3D MEA have comparable SNR and spike statistics to 2D MEAs.

**Fig. 3.**
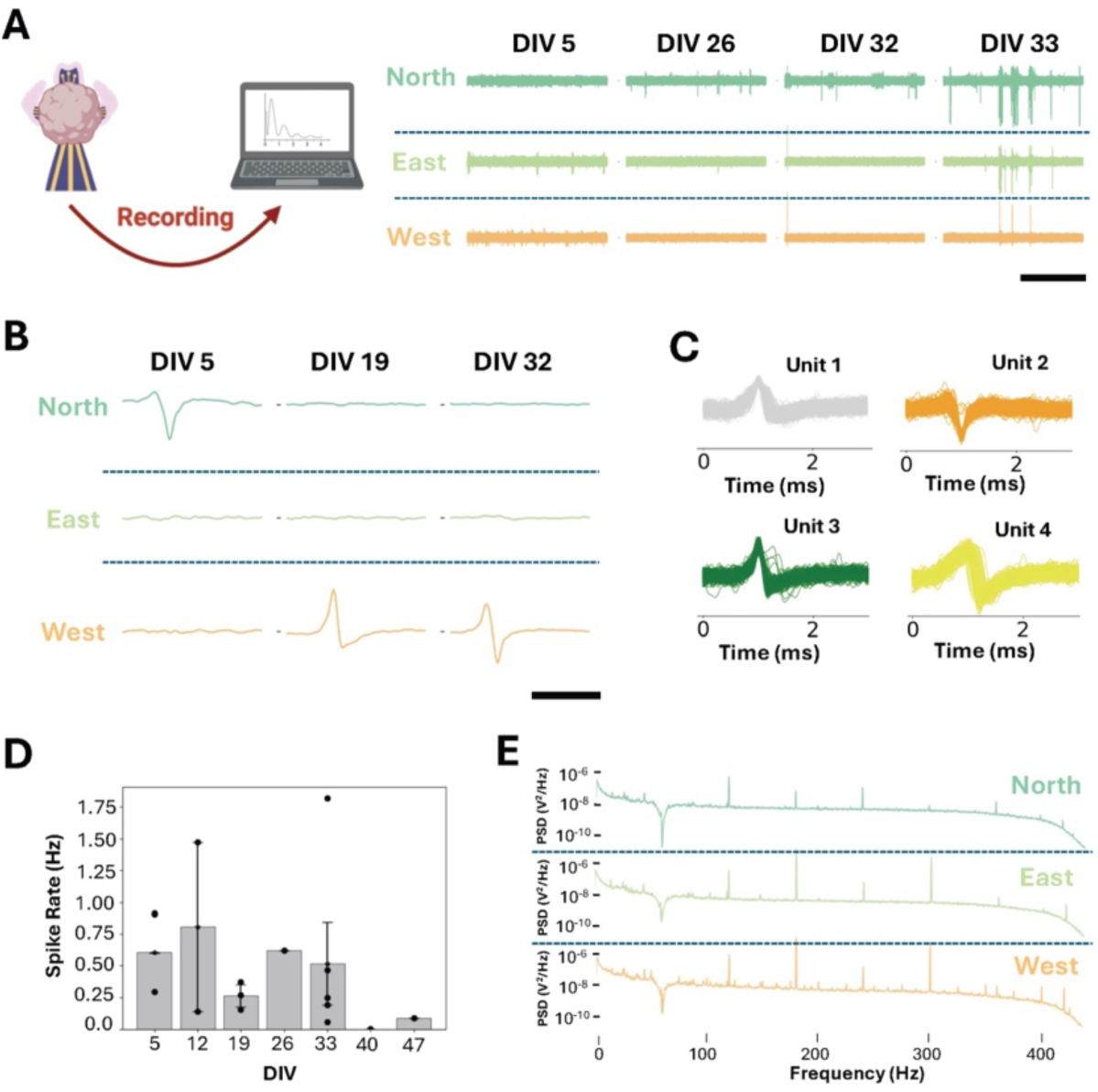
Representative recording traces from NOs measured using three shell MEAs. **(A)** Electrophysiological recording of spontaneous NO neuronal activity. We self-folded shell MEAs on NO differentiated at six weeks and continuously monitored their activity. Observable spiking and burst activity increased throughout the 33 days *in vitro* (DIV) within the shell MEA. Scale bar, 12 sec. **(B)** Analysis of spike differentiation at specific days *in vitro* (DIV). Scale bar, 2 ms **(C)** Four distinct neuronal waveforms (units) were captured throughout all firing. **(D)** Average spike rate frequency displays an initial increase followed by a decrease until around the sixth week. **(E)** PSD analysis of shell MEAs using LFP measurements. The spikes are 60 Hz, and the harmonics are noise.

We also measured regular and consistent signals and observed robust firing up to about 33 days *in vitro* (DIV) with the shell MEA, allowing us to characterize spike rate and interspike interval (ISI) across time and 3D position (**Fig. 3**). After that, signals became too noisy to be sorted accurately. The ISI (**Fig. S12**) shows the temporally recorded signals from three electrode MEAs demonstrating varying spiking activity on the three electrode channels over multiple weeks. Importantly, we observed spatial variations in the single unit statistics across N, S, and W electrodes, highlighting the importance of 3D MEA platforms for monitoring electrical activity in NOs (**Fig. 3A-B**). We filtered the data from each recording channel and noted an amplitude variation at different NO regions, electrode locations (N, E, and W), and DIV. After spike sorting (**Fig. 3B)**, we identified single units per 3D MEA channel with the spike templates, predominately on one spatial channel. Such single-unit response represents the activity of an individual cell as described in **Fig**. **3C** (*29*), indicating that the 3D MEA platform can be used to record individual neurons spatially separated over the surface of the NO.

Using the MountainSort5 algorithm, we clustered and plotted four example units extracted from the 3D MEA signal (Units 1 to 4), plotting each example waveform to demonstrate consistent clustering by spike shape (**Fig. 3C and S12**). To visualize the spike shapes, we plotted the spike trace from the 3D MEA channel with the largest absolute amplitude. The consistent shapes consist of only positive peaks, biphasic waveforms, and negative ones. They are consistent with recordings from individual neurons, including the range of waveform shapes typically observed in extracellular spike recording (*30*). We also plotted the spike rate over DIV. We observed relatively robust spiking over multiple weeks in culture (**Fig. 3D**). The spike rate is consistent with 2D MEA data from similar NOs and falls off after time (*24*). Critically, this spiking demonstrates the ability of the 3D MEA to record single units from across the spatial extent of the 3D NO over days and even weeks. The low-frequency Power Spectral Density (PSD) can also be plotted at different electrodes. We estimated the PSD with the SciPy package using the Welch method. Outside of the peaks, which are harmonics of 60 Hz noise, we observed typical LFP data, including some more prominent peaks, for instance, on the West electrode (**Fig. 3E**). Our findings motivate the spatial analysis of LFP signals to determine the relationships between neural populations in NOs over time.

### 3D Neuromodulation in NOs

To achieve 3D neuromodulation in NOs, we implemented (i) a system that can provide electrical biphasic pulse inputs to the various spatial locations along the surface of the organoid, (ii) a set-up capable of conducting long-term continuous recording, and (iii) an electrophysiology analysis pipeline **(Fig. 4**). Significantly, our MEA design can simultaneously stimulate and record independently at spatial regions all around the contour of the NOs **(Fig. 4A-D**). Confocal fluorescence imaging confirmed the homogeneity of NOs by staining cell nuclei (Hoechst), neuronal stem cells (SOX2), and neuronal axons (NF-H) **(Fig. 4E).** In addition, we imaged mature neurons (NeuN), early-stage neurons (B-III-Tubulin), and neuronal dendrites (MAP2) **(Fig. S10).** The even distribution of neurons, dendrites, and axons throughout the NOs demonstrates that stimulation at any region could spread, enabling responses to be detected by the remaining electrodes (**Fig. 4E and S10**).

**Fig. 4.**
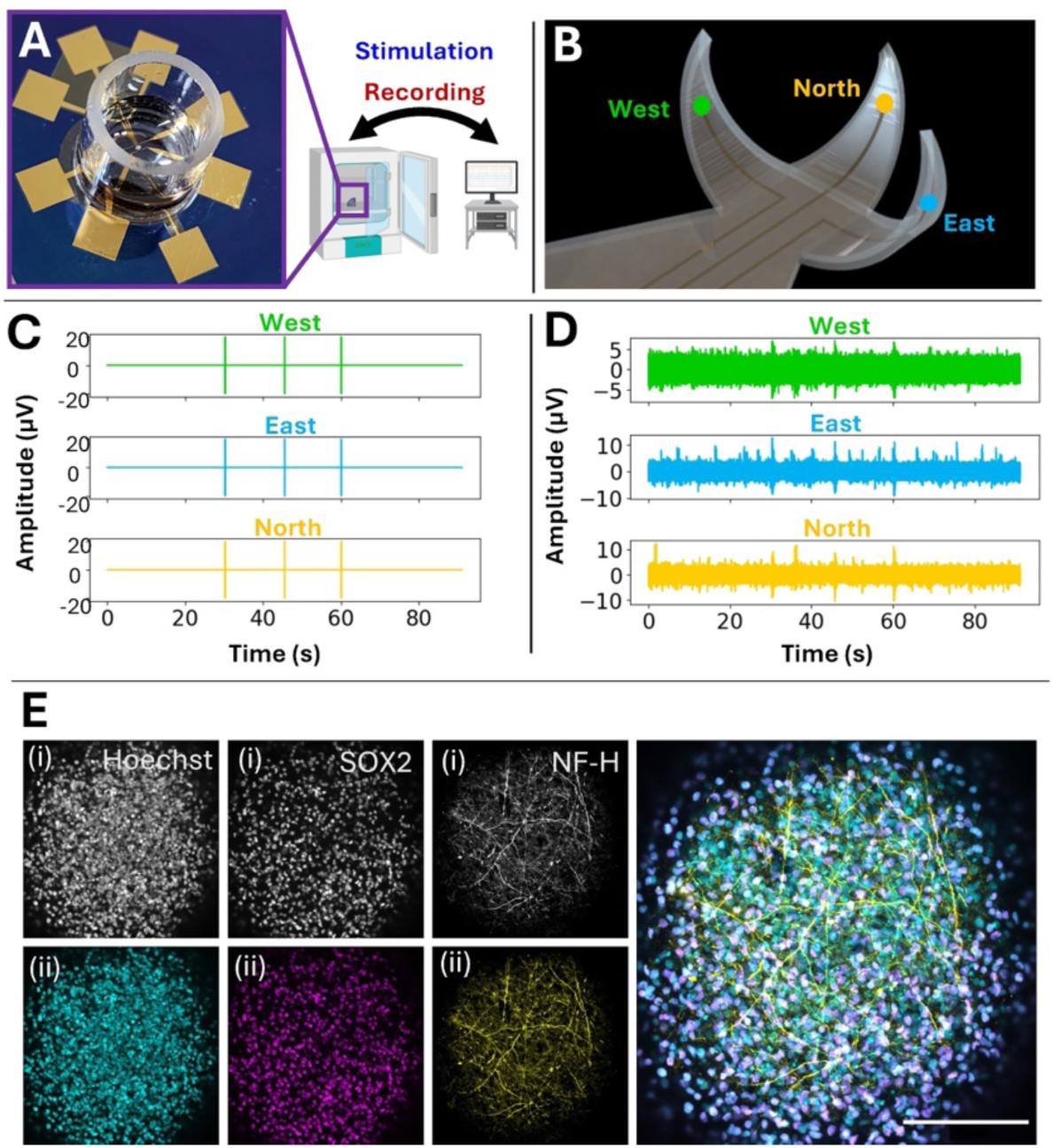
Stimulation recording and using shell MEAs. **(A)** Image of a 3D-shell MEA with an encapsulated organoid. The organoid is stimulated and recorded while MEAs remain inside the incubator. **(B)** Schematic representation of 3D-shell MEA with three leaflets labeled North, East, and West. **(C)** Plot showing the recorded voltage profile when a stimulation current of 20 µA is sent to the organoid through all three electrodes. **(D)** A plot of the recorded voltage profile from all three electrodes shows signals collected from the organoid. The peaks corresponding to the stimulation shown in **(C)** are removed from this profile. **(E)** Representative maximum intensity z-projection images of eight-week-old organoids show the presence of nuclei (Hoechst), neuronal stem cells (SOX2), and axons (NF-H) shown in grayscale (i) and turquoise, purple, and yellow respectively (ii). The staining illustrates cellular homogeneity within the organoid. Images were taken at 20x. The scale bar is 100 µm.

We adopted parameters based on 2D MEA NO experiments (*2*) for 3D neuromodulation using the shell MEAs. We delivered stimulation parameters via a cathodic-first, biphasic pulse of 233.3 μs (**Fig. 5A**). We applied such pulses with amplitudes ranging from 5 μA to 60 μA and concurrently recorded the activity and observed an increased spike rate following a 20 – 30 µA stimulation (**Fig. S13**). As indicated by the CIC studies, our self-folded shell MEAs with PEDOT: PSS coating can provide these stimulation pulses without causing water electrolysis. Remarkably, we observed that spiking activity increased with peak stimulation currents of 20-30 µA (**Fig. 5B, C)**, suggesting that this stimulation can successfully drive the activity of individual neurons at different spatial locations, which returned to baseline firing levels after approximately 10 seconds (**Fig. 5C**).

**Fig. 5.**
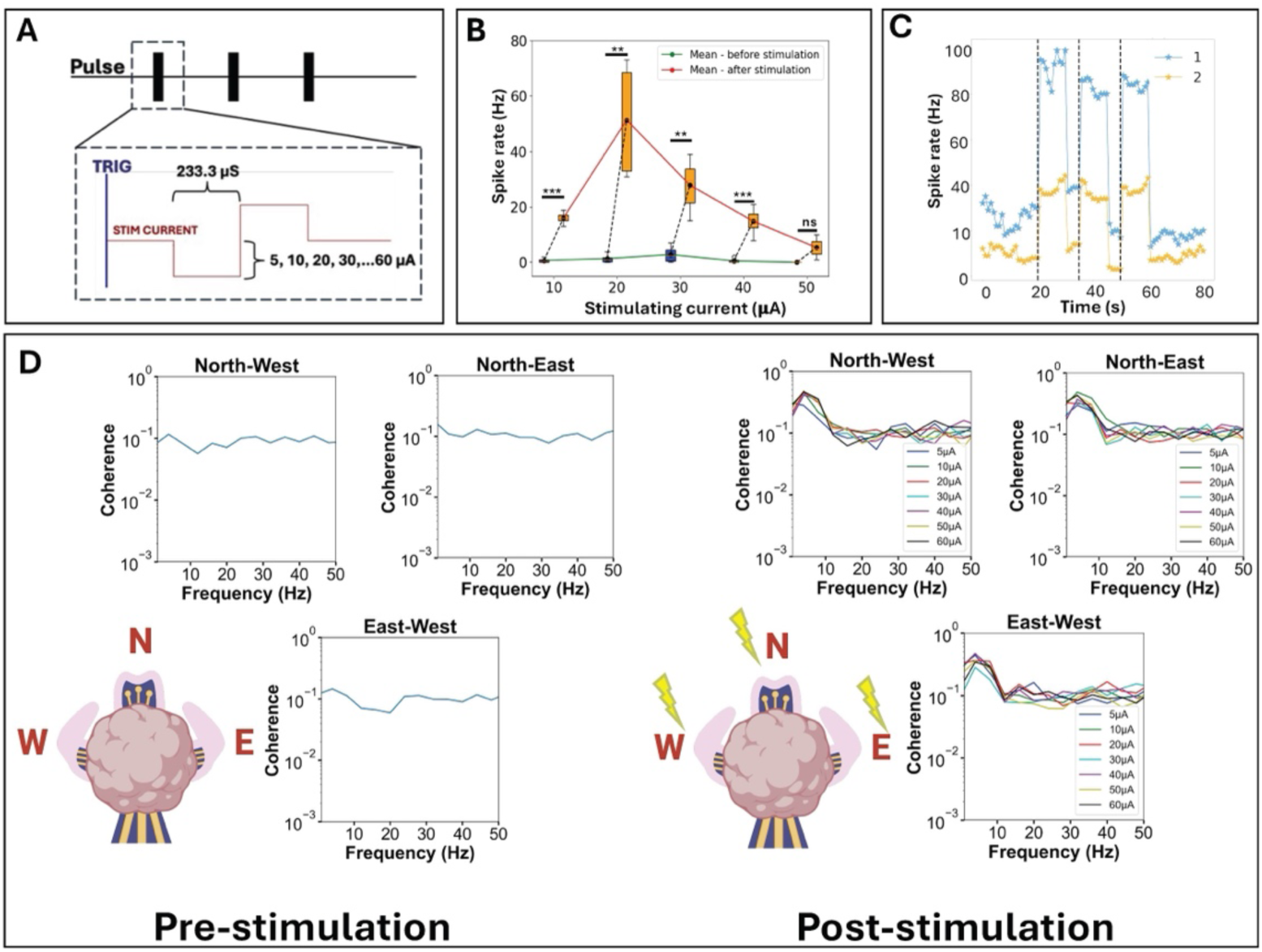
Neural response and coherence analysis under varying stimulation intensities. **(A)** Stimulation Protocol: (i) Schematic of pulse train used for stimulation. (ii) Zoomed in view of a single stimulation pulse with 233.3 µs duration and current intensities from 5 to 60 µA. **(B)** Bar graph showing detected spikes before and after stimulations across different current intensities. Significant changes are indicated with statistical markers (p<0.01, p<0.001). **(C)** Firing activity of two detected units (1 and 2) over a 10-second time window during stimulation at 20 µA. Both neurons show increased activity upon stimulation (indicated by the dotted lines). **(D)** Pre-stimulation coherence spectra show baseline coherence across different electrode pairings. Post-stimulation coherence spectra illustrate changes in coherence for varying stimulation intensities. Inset diagram shows electrode positions with respect to the NO (N, E, W directions).

We repeated stimulus levels thrice for a given NO for each stimulation level. **Fig. 5B** shows the change in spike rate, averaged over units recorded from three electrodes, for the different stimulation levels. Error bars show the standard deviation for each stimulus level. Significance testing indicates that the stimulation can repeatedly and reliably drive spiking activity in the 3D organoid, with a preferred peak of 20-30 μA. **Fig. 5C** shows the spiking rate for two different units over a recording session of three stimulation pulses. Individual data points represent an estimate from a short time window. For both units, activity rapidly increases after stimulation and then decays back to baseline levels. Stimulation consistently drives spiking activity from individual neurons using the 3D MEA in a targeted fashion.

### Stimulation effects of LFP and NO coherence analysis

To further characterize the spatiotemporal dynamics of neuronal responses, we analyzed the coherence changes between electrode pairs before and after stimulation with different current intensities (**Fig. 5D**). LFPs are low-frequency components of extracellular potential that mainly reflect the synaptic activity of neuronal populations within a localized region (*31–34*). We analyzed the LFPs using coherence analysis (from 0 to 1) to assess the synchronization of neural signals between different areas of NOs (**Note S2)** (*33*, *35–39*).

We first analyzed spontaneous LFP activity across various regions of the NO. Analysis of 5-minute NO recordings from our NOs revealed consistently low coherence values (around 0.02) between electrode pairs in the 1-50 Hz pre-stimulation range (**Fig. S14**). This pre-stimulation low synchronization suggests that different NO regions maintain relative independence in their activity patterns. We then analyzed the coherence of LFP activity from stimulated NOs. Given that the neuronal firing rate peaks within 0.25 s after stimulation (*2*), we used a window of 0.25 s to capture the frequency changes more efficiently. We performed the same processing that was used on the pre-stimulation signal. The results showed that the coherence values of all electrode pairs pre-stimulation remained around 0.1 for different stimulation intensities in the 10-50 Hz frequency range (**Fig. 5D**). In the lower frequency range (1-10 Hz), we detected a significant coherence of about 0.5 immediately after stimulation, accompanied by substantial power enhancement in the PSD analysis (**Fig. 5D and S15**). As a control, stimulation of a silver bead (positive control) showed high coherence values (above 0.5) between all electrode pairs, confirming that signals from a simple conductor maintain intense phase relationships regardless of spatial location (**Fig. S16**). The coherence in NOs may involve neuronal propagation, where the resulting waves synchronize activity between different spatial areas and cross-region communication (*40*, *41*). We note that since our design uses a single-wire reference, we cannot entirely rule out contributions to coherence at low frequencies that may arise from volume conduction effects (*36*). Overall, the LFP signals between the three electrodes maintained coherence immediately after stimulation but low coherence before and well after (>2.5 s) stimulation, demonstrating that stimulating NOs recover and that different spatial regions generate distinct signals. Subsequent 3D thermograms will provide more evidence of the changes in the firing rates of neurons from various locations.

### 3D Spatiotemporal signal modeling and modulation

We analyzed recordings from three and 16-electrode 3D MEAs and developed a method for 3D visualization and modeling of NO responses. As described in the Methods section, we mapped the activity from each electrode to the 3D sphere corresponding to the approximate spatial extent of the NO. We estimated the positions of the folded electrodes and mapped the activity to each electrode. We can map LFP amplitudes or per-channel spike events (multi-unit activity, the processing pipeline results before spike sorting) to the 3D MEA. We then spatially smoothed and plotted the activity in 3D over time. **Fig. 6** shows the results of this approach utilizing multi-unit activity (MUA) from the 16-electrode MEA in response to stimulation. **Figs. 6A** and **6B** show the position of the 16 electrodes relative to the spherical organoid before folding (in a 2D configuration) and the estimated position after folding. Similar to the response curves in **Fig. 5B**, **Fig. 6C** shows the change in spike rate from identified single units in response to repeated stimulation with different current levels. Again, we observed consistently elevated activity levels. Using per-channel MUA, we demonstrated that we can create an event raster per channel (**Fig. 6D**) and map activity to a spherical model of the NO (**Fig. 6E)**. We can see the spatial extent of activity over the surface of the organoid. Moreover, we can see responses to stimulation pulses over time, including activity propagation through the NO. This approach allows for comparative visualization of 3D NO activity over time and quantifying spatial activity in response to stimulation, creating a visualization of spatiotemporal neuromodulation (**Fig. 6F and Movie S1)**.

**Fig. 6.**
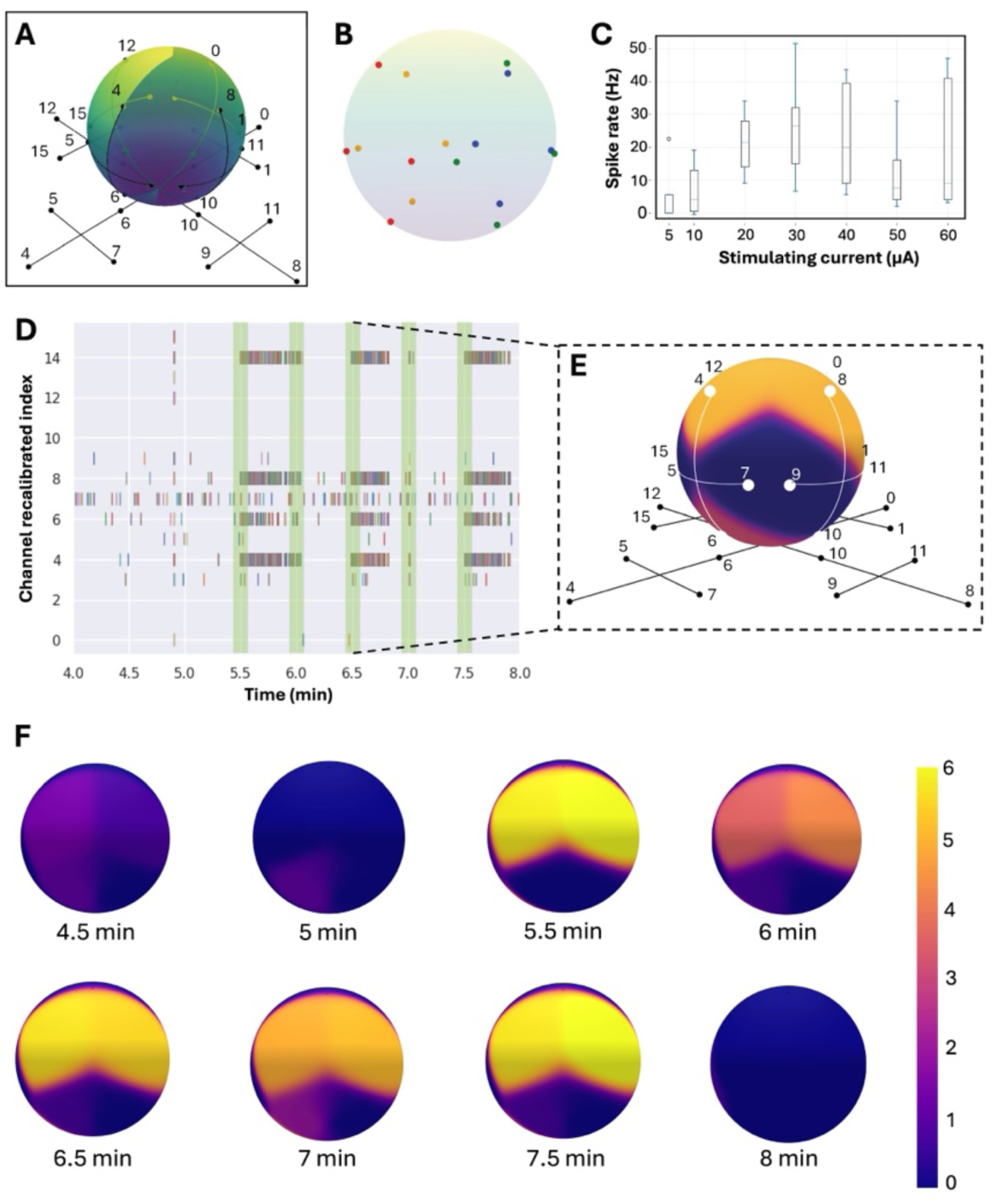
Spatiotemporal neuromodulation. **(A)** Schematic representation of numbered 16-shell MEA with encapsulated NO. The positions of the electrodes are shown before and after self-folding. **(B)** Unit sphere mapping of the 16 electrodes where each color represents a leaflet, calculated geometrically assuming a spherical NO. **(C)** The average change in spike rate for 16-shell MEA electrical stimulation, which is averaged over all identified units, using a time window before and after the stimulation, following the procedure for Fig. 5. **(D)** Raster plot of 30 μA stimulation pulse given at 30-second intervals, with each row representing an electrode and each tick a detected event (MUA) and the shaded areas representing the stimulus period (green). **(E)** 3D spatially smoothed log+1 exceedance counts plot of evoked multi-unit response in NO, showing activity distribution over the three-dimensional sphere. **(F)** Images of spatiotemporal neuromodulation visualized using the multi-unit evoked response at eight-time points during the stimulation, demonstrating neuromodulatory changes in the MUA over space and time.

## DISCUSSION

Neuromodulation is a key step in realizing all biocomputing and organoid intelligence (OI) systems. Also, due to the small size and 3D shape of NOs, there is a need to create biocompatible electronic microsystems capable of stable and long term interfacing in 3D. We designed and fabricated the 3D shell MEAs with careful attention to the use of biocompatible materials, soft conductive electrode pads, multiple distributed electrodes around a sphere, and electronic control systems that enable simultaneous stimulation and recording. Notably, the fabrication processes are scalable, cost-effective, and amenable to reliable CAD-based photolithography and strain engineering. No transfer steps between substrates are needed, and shells can be fabricated on silicon or glass substrates for integration with ancillary microelectronic or microfluidic modules. We note that due to cost limitations, we utilized a relatively inexpensive mask set with the smallest resolved feature size of 20 microns, allowing the fabrication of 3-electrode and 16-electrode shells. Importantly, our recordings observed similar stimulation and recording trends in three-electrodes and 16-electrode shell MEAs, as shown in **Fig. 5** and **Fig. 6**, respectively. A more significant number of electrodes, wires, and contact pads are achievable with higher-resolved masks using the same process, albeit at a higher cost.

We monitored the electrical activity of the NOs inside shell MEAs for several weeks. In a controlled, isolated environment, we demonstrated that we can simultaneously measure and stimulate a single NO at 3D spatially varying electrode points. Using a spike-sorting algorithm, we could detect single units at high temporal resolution at a high induced activity from 20 – 30 µA. Stimulation and recording data from the proposed platform could help us understand how to design neuromorphic and non-von Neumann computational architectures and provide an advanced model of neural development and neurological disorders. In the future, we seek to investigate 3D spatiotemporal patterns and incorporate neurochemical stimulation in response to diverse chemical and electrical stimuli and to model closed-loop feedback learning paradigms. We observed neuromodulation trends in activity in both three-electrode and 16-electrode 3D MEAs, with prominent changes in spike activity in response to stimulation. Further increasing electrode density with the 3D MEA design will enable greater insight into the development of spatial responses in NOs and improve the ability to encode stimulation information and activity states from NOs.

Critically, our results showed the ability to record spatially isolated single units and LFP signals simultaneously. Both signals showed modulation in responses due to stimulation, including increased coherence. The 3D MEA, therefore, allows for the simultaneous recording of 3D spatiotemporal activity between spatially isolated signals (individual neurons) and spatially coupled signals (LFPs). Moreover, our 3D visualization approaches allow for the spatial representation of multi-unit and LFP activity in spontaneous and stimulated conditions. In the future, spike rates can be used to monitor 3D spatiotemporal activity for experimental conditions such as differentiating vs. mature NOs, different disease states, and more. Spatiotemporal variations in stimuli vs. response can yield insight into the types of neuronal self-organizing networks. Also, we can leverage this approach to study the fundamental neuroscience of plasticity and population activity in NOs, characterize the responses of different NO models (for instance, different disease states), and even form the basis of bioprocessors.

## MATERIALS AND METHODS

### Fabrication of the shell MEAs

We fabricated shell MEAs following a previously reported process flow with some modifications (*14*). We utilized thermal oxide (300-500 nm) coated silicon wafers as substrates, designed all photomasks using AutoCAD, Autodesk, USA, ordered all masks from Artnet Pro Inc, San Jose, CA, obtained all photoresists from Kayaku, Westborough, MA and carried out photopatterning using an NXQ-4000 mask aligner with a mercury arc lamp (Neutronix-Quintel, Morgan Hill, CA). The typical fabrication involved seven photomasks for patterning a sacrificial layer followed by differentially crosslinked SU8 layers, wiring, electrodes, contact pads and bond pads. We first patterned 50 nm germanium (Ge) sacrificial layer on the wafers lift-off metallization. Briefly, we spin-coated Shipley SC 1827 photoresist at 3000 rpm and then baked it at 115°C for 1.5 minutes to obtain ∼2.7 μm film. We exposed the coated wafer through a photomask (150 mJ/cm^2^) and developed the photoresist for one minute in a diluted Microposit 351 developer (one part developer and five parts deionized water). We thermally evaporated 50 nm of Ge (Kurt J. Lesker Company, Jefferson Hills, PA) and then dissolved the resist in acetone. Next, we patterned the SU8 layers of the self-folding shell by first spin-coating SU8 2005 (Kayaku, Westborough, MA) at 3000 rpm as the first SU8 layer (∼5 μm thickness) and soft-baked at 65°C, 95°C, and 65°C for 1 minute, 3 minutes, and 1 minute, respectively. We exposed the first layer to an energy density of 240 mJ/cm^2^ for our three-electrode shell and 160 mJ/cm^2^ for our 16-electrode shell MEA design through a photomask, achieving a patterned crosslinking of this SU8 layer. We post-exposure baked the wafer at 65°C, 95°C, and 65°C for 1 minute, 3 minutes, and 1 minute, respectively. Subsequently, we developed the wafers in SU-8 Developer (Kayaku, Westborough, MA) for 1.5 minutes to remove the uncrosslinked SU8.

Next, we patterned a gold (Au) electrode layer and a copper (Cu)-covered peripheral bond pad on the first SU8 layer using a lift-off patterning process like the Ge patterning step. We used 10 nm chromium (Cr) to ensure good adhesion between the Au layer (50 nm) and the SU8 layer. Unlike in the case of Ge, we carried out the lift-off in isopropyl alcohol (IPA), as acetone affects the first SU8 layer and, consequently, the self-folding behavior. We left the wafer inside IPA for 24 hours and then sonicated it to remove the SC 1827 photoresist. To improve the success rate of wire bonding, we deposited an additional 150 nm of Cu at the bonding pads.

We selectively insulated the patterned Au layer everywhere except the leaflets and contact pad electrodes by coating and patterning a second layer of SU8 2005 on the patterned electrode layer, following an identical process delineated earlier for the first SU8 layer. We then patterned the leaflets with a partially crosslinked third SU8 layer, forming a differentially crosslinked bilayer with self-folding capability. We partially crosslinked this layer by exposing the region to a lower energy density than the first layer (120 mJ/cm^2^). All the other steps remained the same as the first SU8 layer patterning. Additional details and a more detailed process flow are in the SI.

### NO integrated packaging

We designed the packaging for the shell MEAs (with an 18-pin connector adaptor) using the open-source KiCAD software (KiCad Services Corporation, Long Beach, CA). We used this design to order an Electro-less Nickel /Immersion Gold (ENIG) surface finish printed circuit board (single PCB, 1.6 mm PCB thickness, JLCPCB, Hong Kong, China). The wafer with shell MEA patterns was diced using a diamond tip scribe to a desired 25 mm x 25 mm piece and bonded to the integrated package using polydimethylsiloxane (PDMS) mixture (10 parts monomer and 1 part curing agent) at 85°C for 15 minutes. We wire-bonded the peripheral bond pads on the diced shell MEAs to the bond pads on the PCB board with a 25 μm diameter gold wire using K&S 4526 Manual Bonder (Kulicke and Soffa Industries, Fort Washington, PA). For wire bonding, we heated the substrate to 95°C to prime the bond pads and PCB for bonding. Following the recommended machine adjustment parameter (*42*), our wire feed was at 30°/45°, and the power, contact time, and force to bond the wire to the peripheral bond pads on the integrated packaging were at 1.8 W, 4.5 ms, and 6.5 g, respectively. We set our second bonding point to the MEA electrode patches to a power of 2 W, time of 4 ms, and force of 1 g. We soldered the two 9-pin connectors with a 2.54 mm pitch to the Neural organoid-integrated packaging to allow for easy interfacing to obtain external electrophysiology readings. Furthermore, we isolated the region with the reference electrode and the self-folding leaflets with contact pads from the wire-bonded electrode patches by bonding a Corning™ PYREX™ cloning cylinder (Fisher Scientific, Hampton, NH) of diameter 10 mm and height 10 mm to the region using PDMS (**Fig. S6**).

### PEDOT: PSS wlectrode coating and characterization

We prepared solutions for electrodeposition of PEDOT: PSS by mixing 10 mM EDOT (Sigma-Aldrich, St. Louis, MO) and 0.4 wt % of PSS (Sigma-Aldrich, St. Louis, MO) in DI water. We pipetted this solution into the cloning cylinder and made connections to the working electrodes using alligator or pogo pin connector clips. We used a platinum (Pt) rod as the counter electrode to electrodeposit 30 μm radius contact electrodes with a current density of 3.96 mA/cm^2^ for 4 minutes to create a 10 μm thick coating. We applied the same current density to the reference electrodes with a radius of 75 μm and deposited the film for 6 min. After electrodeposition, we rinsed the wafer with DI water.

We characterized our electrode impedance using Electrochemical Impedance Spectroscopy (EIS) with the VersaSTAT3 Potentiostat Galvanostat (AMETEK Scientific Instruments, Oak Ridge, TN). For this characterization, we used a three-electrode set-up involving the working electrode, an Ag/AgCl reference electrode, a Pt counter electrode, and 1x Phosphate-Buffered Saline (PBS). We applied a frequency sweep from 1 Hz to 10 kHz on bare and PEDOT: PSS-coated gold electrodes (n=3, **Fig. S7**). Using the same set-up, we measured the charge storage capacities (CSCs) of the electrodes by performing cyclic voltammetry on both the bare and the PEDOT: PSS-coated gold electrodes from –0.6 V to 0.6 V range, which is the intersection of electrolysis windows of gold and PEDOT: PSS, at a scan rate of 200 mV/s (*23*) (**Fig. S8**). Furthermore, we characterized the voltage transient of the stimulation pulses to determine the charge injection capacity (CIC) by checking if the polarization potential exceeds the electrolysis potential.

### NO generation protocol

As previously published, NOs were generated following our in-house protocol (9,26). We purchased female fibroblast-derived NIBSC8 (N8) iPSCs from the National Institute for Biological Standards and Control (NIBSC, UK), complete with a certificate confirming the absence of mycoplasma, bacteria, or viruses and a normal karyotype verified by SNP Array. We cultured these hiPSCs in mTESR-Plus medium (StemCell Technologies, Vancouver, CA) under 5% O_2_, 5% CO_2_, and 37°C. We validated the stemness of the cells using immunocytochemistry and flow cytometry for Oct4, Nanog, TRA-1-61, and Sox2 markers (*25*). Subsequently, the hiPSCs were differentiated into neural progenitor cells (NPCs) in a monolayer using a serum-free neural induction medium (Gibco, Thermo Fisher Scientific). We expanded Nestin/Sox2-positive NPCs, dissociated them into a single cell suspension, and seeded them into uncoated 6-well plates. These 6-well plates were placed on a shaker with continuous gyratory shaking (80 rpm, 19 mm orbit) at 37°C, 5% O_2_, and 20% CO_2_ to form NOs. After 48 hours, we induced differentiation by substituting the neural induction medium with neural differentiation medium comprising the B-27™ plus kit, 1% Glutamax (Gibco, Thermo Fisher Scientific), 10 ng/ml human recombinant GDNF (GeminiBio™), 10 ng/ml human recombinant BDNF (GeminiBio™), and 1% Pen/Strep/Glutamine (Gibco, Thermo Fisher Scientific). We changed the neural differentiation media every two to three days during recordings.

### NO immunocytochemistry

We conducted immunocytochemistry following the outlined protocol (*25*). We fixed the organoids with 2% PFA, permeabilized them using 0.1% Triton X, and blocked them with 100% BlockAid™. We applied primary antibodies (**Table S1**) at 4°C on a shaker for 24 hours and then stained them with secondary antibodies under the same conditions for 24 hours. We used Hoechst 33342 trihydrochloride (Invitrogen Molecular Probes) to stain nuclei at a 1:10,000 dilution. We performed imaging using an FV3000RS Olympus confocal microscope.

### NO shell MEA encapsulation

We dissolved the Ge sacrificial layer in a 6% hydrogen peroxide solution at room temperature for 24 hours to selectively detach the leaflets from the oxide-coated silicon substrate (*14*). Afterward, we immersed our SU8 electrode bilayer in acetone for five minutes as a preconditioning step to remove all non-cross-linked SU8 monomers to facilitate our folding actuation and sterilize the glass chamber. We aspirated acetone from the cylindrical chamber and flushed thrice with 1x PBS solution to remove the acetone residue. This process initiates self-folding through the solvent exchange (*43*). Finally, we transferred 300 μl of fresh neural differentiation culture media containing an organoid, moved it to the center of the self-folding leaflets, and incubated it at 37°C for 1 hour to complete encapsulation. To ensure organoid viability, we aspirated 150 μl of liquid every two days and replaced it with fresh media.

### Electrophysiology data acquisition

We acquired electrophysiology data using an Intan RHS Stim/Recording System (Intan Technologies, Los Angeles, CA). We acquired data through the RHX Software interface, with recording channels sampled at 30 kHz, an amplifier bandwidth of 1.17 Hz-7.60 kHz, and a 60 Hz notch filter. We converted the raw data into the Neurodata Without Borders (NWB) format, adding key experimental metadata to the files. After conversion, data were analyzed in Python using the SpikeInterface package (*27*). We controlled precise stimulation times and amplitudes with the Intan RHS amplifier, specifying stimulation time, pulse width, and amplitude (in amperes). Full details of the process are described in **Note S1**.

### NO signal processing

After acquiring data, we processed the files in Python using the SpikeInterface package (full details in **Note S1**). We bandpass filtered each channel between 300 and 3000 Hz for spike sorting, followed by channel whitening. We detected putative neuronal events within a default threshold of 5.5 times the standard deviation of the signal and used the MountainSort algorithm (*28*) to cluster spikes based on Principle Component Analysis features. Two analysts manually inspected spike clusters and template waveforms, identifying clusters that required merging and clusters that contained only noise. We logged spike times and spike waveforms for further analysis. We identified single units for each recording to compute spike rates and inter-spike interval histograms. We used artifact rejection to detect time windows with an amplitude greater than 500 mVs to assess stimulation-based modulation. We omitted these areas from the analysis and used a window of 0.5 seconds before and after the detected artifacts to compute baseline and stimulus-driven activity in each recording (**Fig. S12 and S13**).

We generated LFP and MUA signals by bandpass filtering the raw signal between 1-400 Hz and 300-6000 Hz (the same range used for spike sorting). We computed spectrograms and power-spectral density estimates (using Welch’s method) using the Python package SciPy (*44*). For stimulation experiments, we stored signal sequences on an Arduino microcontroller. We used them as a trigger signal for the Intan amplifier, ensuring sub-millisecond precision of the stimulus pulse train.

For the 3D visualization, we analyzed windowed pre/post-stimulus MUA and multi-unit channel-level statistics to estimate the spatial effects of stimulation, utilizing the output of the event detection step. Given a presumed organoid radius and detected events on each channel, we folded each electrode in idealized unit spherical arcs around each organoid. Next, we smoothed it using a Gaussian kernel with distance computed along the sphere. **Fig. 6** represents the smoothing where the value is the log of the post-stimulus exceedance counts in a five-second window with the 16-electrode shell system. Additional details are in **Note S3**.

## Acknowledgments

We acknowledge Dr. Brett Collar for assistance with wire bonding and Dr. Ashlee Liao for discussions. We also thank Soo Jin Choi and Zhaoyu Liu for their discussions in developing the code used for 3D spatial mapping. This publication was developed under Assistance Agreement No RD83950501, which was awarded by the U.S. Environmental Protection Agency to Drs. Smirnova and Gracias. It has not been formally reviewed by the EPA. EPA does not endorse any products or commercial services mentioned in this publication. Dowlette-Mary Alam El Din was supported by the National Institutes of Health (T32 ES007141) and the International Foundation for Ethical Research Graduate Fellowship. The authors also acknowledge funding from the Johns Hopkins University SURPASS Program and the National Science Foundation (NSF) EFMA-2318093 and CBET-2348680. The views expressed in this document are solely those of the authors and do not necessarily reflect those of the funding agencies (EPA, NIH, and NSF).

## Author contributions

Conceptualization: CA, DG, ECJ, LS, DHG

Neural organoids generation: DMA, LS

Device fabrication: CA, DG, HS, ZO, EL, GMM, PRR

Electrophysiological measurements: CA, DG, HS

Optical Imaging: CA, DG, DMA

Data analysis: ECJ, LCD, BSC, BM, YL, ZO, HS

Supervision: LS, ECJ, DHG

Images and Illustrations: CA, DG, ZO, HS, EL, GMM, PRR

Original draft manuscript: CA, DG, LCD, DMA, BM, ZO, EL, GMM, PRR, BSC, ECJ, LS, DHG

## Supplementary Materials

Materials and Methods

Supplementary Text

Figs. S1 to S16

Tables S1

References (1–7) Movies S1

