## Supplementary Information for "Neuromodulation in neural organoids with shell MEAs"

### Supplementary Materials for
#### **Neuromodulation in neural organoids with shell MEAs**

**Authors**

Chris Acha[1]†, Derosh George[1]†, Lauren C. Diaz[2], Ziwei Ouyang[3], Dowlette-Mary M. Alam El Din[4], Hrishikesh Surlekar[1], Babak Moghadas[5], Eva Loftus[6], Gandhali M. Mangalvedhekar[1], Pratyush Sai R. Rayasam[3], Yu-Chiao Lai[7], Lena Smirnova[4,8], Brian S. Caffo[5], Erik C. Johnson[2], David H. Gracias[1,8,9,10,11,12,13]*

**This PDF file includes:**

Figs. S1 to S16
Tables S1
Supplemental Note 1 to Note 3
References (1 to 7)

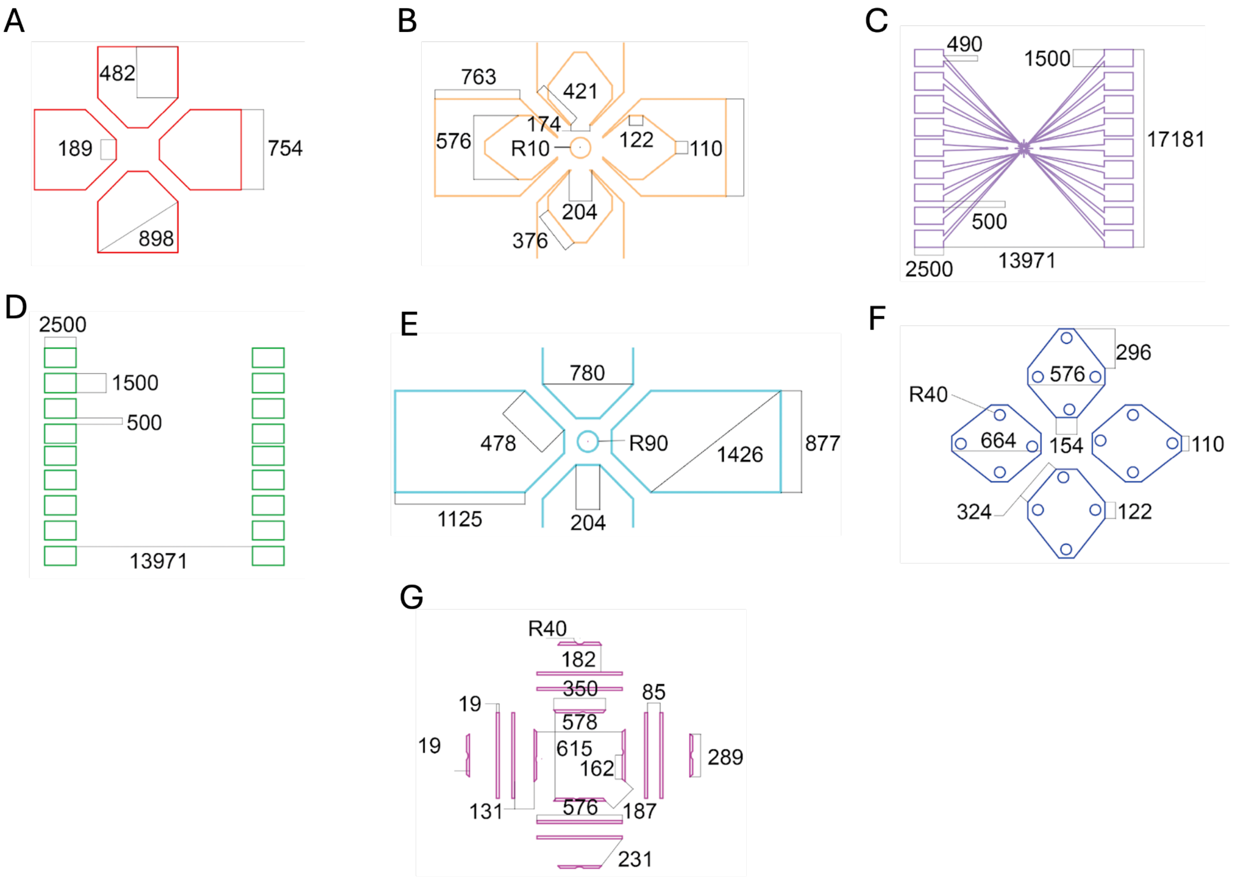


**Fig. S1. Details of CAD layouts used in each of the seven masks used to fabricate 16-electrode 3D Shell MEAs. (A)** Sacrificial Ge layer, **(B)** first SU8 layer, **(C)** Cr/Au electrode layer, **(D)** wiring and bond pad, **(E)** second SU8 layer, **(F)** third SU8 layer, and **(G)** second exposure of the third SU8 layer. All dimensions are in microns, and R refers to the radius.

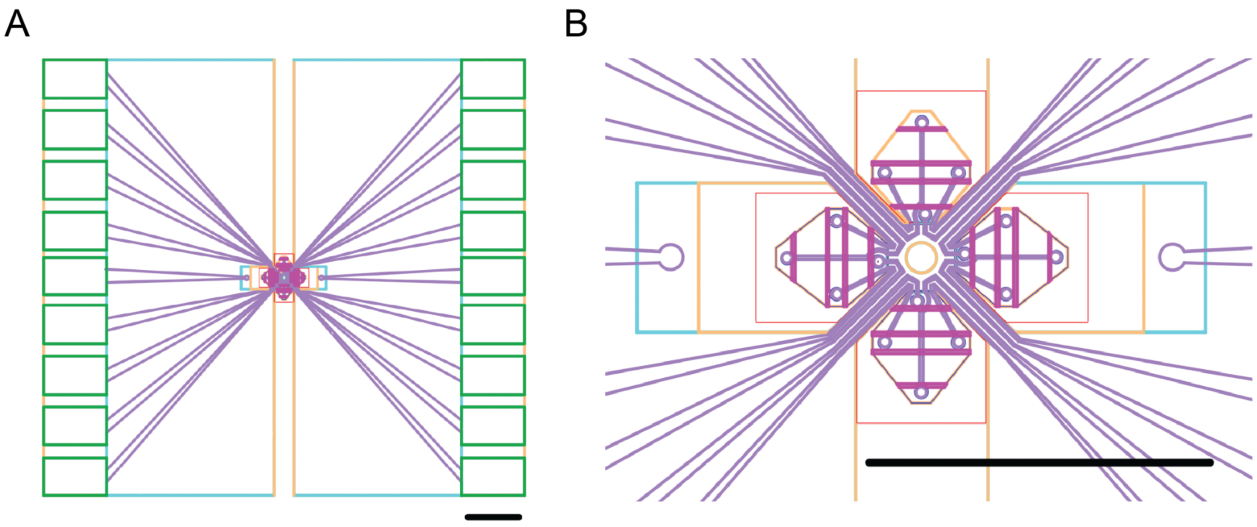


**Fig. S2. Overlay of the CAD masks to fabricate 16-electrode 3D shell MEAs. (A)** Zoomed-out image of the shell with bond pads. **(B)** Zoomed-in regions showing the six CAD mask layers of the four-leaflet shell MEAs. Black lines indicate scale bars and are 2 mm.

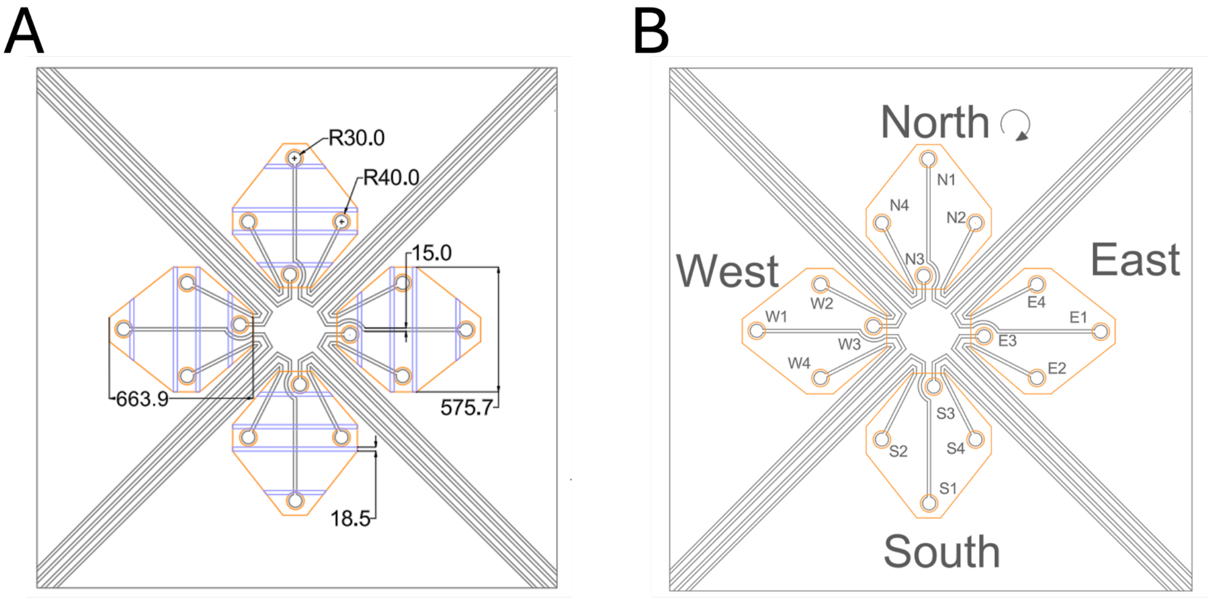


**Fig. S3. CAD mask zoom-in of the electrode numbering scheme used for 16-electrode 3D shell MEA. (A)** CAD mask. **(B)** Cardinal orientation of the electrodes (North, South, East, and West), with individual electrodes numbered in a clockwise sequence starting from tip to base of each petal.

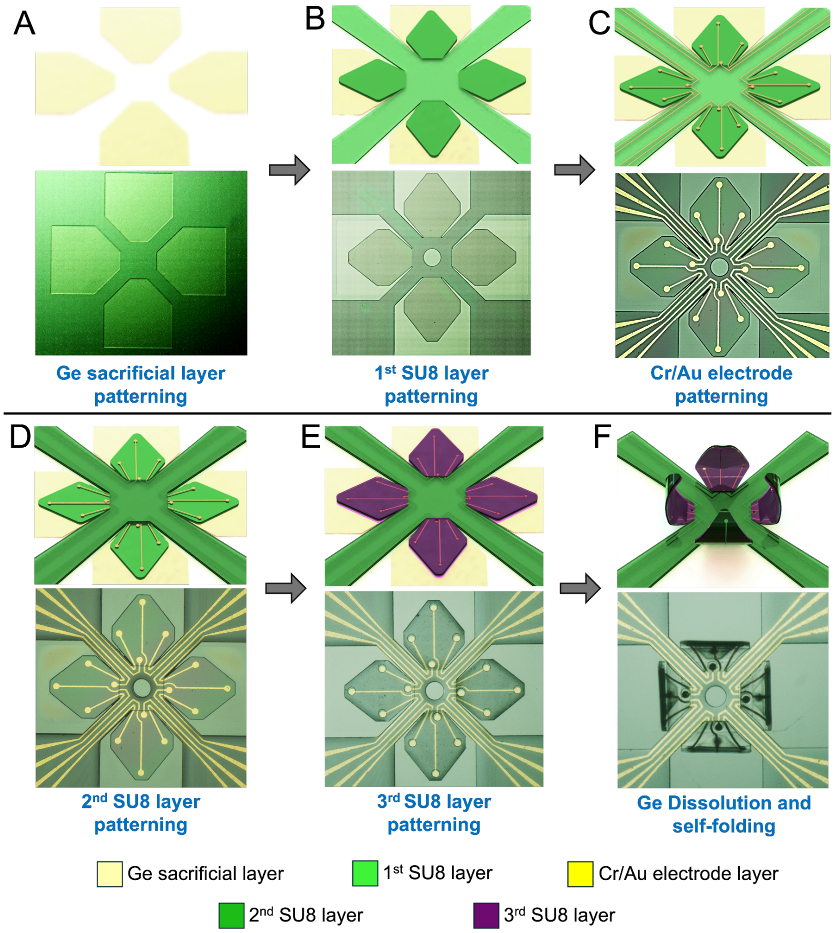


**Fig. S4. Schematics and experimental brightfield microscopy images at different stages during the microfabrication process with the seven CAD masks. (A)** We first evaporated a 50 nm thick germanium (Ge) sacrificial layer onto oxide-coated silicon wafers. This Ge layer allows the subsequent SU8 layers to remain planar and adhere to the substrate during fabrication. We later dissolved the Ge sacrificial layer to release the leaflets for self-folding. The Ge layer was patterned using a CAD mask and a lift-off process in the shape of the shell leaflets with an overhang (see note on CAD mask design for sacrificial layer 1 in Fig. S1). **(B)** We then spin-coated SU8 2005 photoresist at 3000 rpm and selectively UV cross-linked (160 mJ/cm²) using a second photomask. **(C)** A 10 nm layer of chromium was first evaporated onto the patterned SU8 layer to promote strong adhesion between the gold and the SU8 surface. Following this, we evaporated a 50 nm layer of gold to form the conductive pathways for the electrodes. The electrodes extend from the central region, where the organoid will be placed, to the outer bond pads on the wafer for signal measurement. **(D)** A second SU8 2005, approximately 5 μm thick, was spin-coated over the patterned gold electrodes at 3000 rpm and exposed to UV light at 150 mJ/cm² as an electrically insulating layer. **(E)** A third SU8 2005 layer was spin-coated and exposed at a lower intensity of 120 mJ/cm². **(F)** The Ge sacrificial layer was dissolved using 6% hydrogen peroxide at room temperature for 24 hours, releasing the shell leaflets. The wafer was treated with acetone for 5 minutes and then washed thrice with DI water to remove the non-crosslinked SU8 to trigger folding.

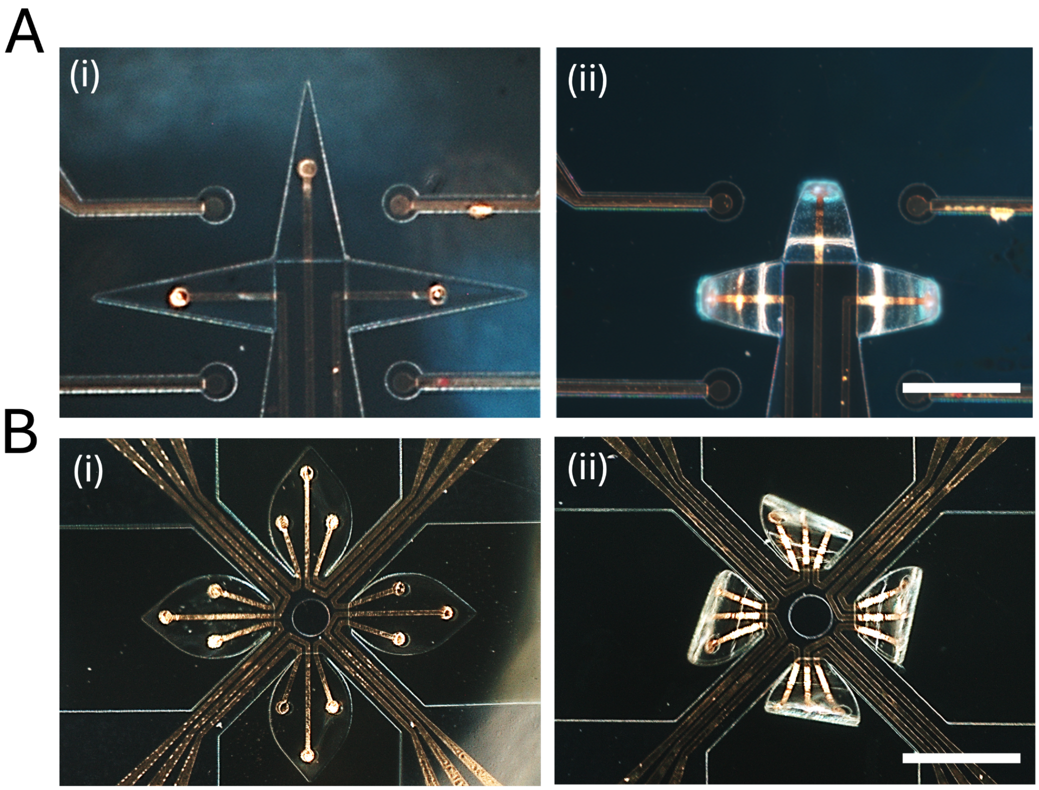


**Fig. S5. Multi-electrode (three- and 12-electrode) shell MEAs pre and post self-folding. (A)** (i) Top view image of three-electrode shell MEA before the Ge sacrificial layer was dissolved with 6% hydrogen peroxide, showing a planar configuration with leaflets flat against the wafer. (ii) Image post-dissolution of the Ge sacrificial layer using 6% hydrogen peroxide freed the leaflets, resulting in self-folding. **(B)** (i) Top view image of the 12-electrode shell MEA before dissolving the Ge sacrificial layer, showing a planar configuration, with electrodes symmetrically positioned across each leaflet. (ii) Top view image post-dissolution of the Ge sacrificial layer using 6% hydrogen peroxide. Upon dissolution, the leaflets were released, resulting in self-folding. Scale bars are 500 µm.

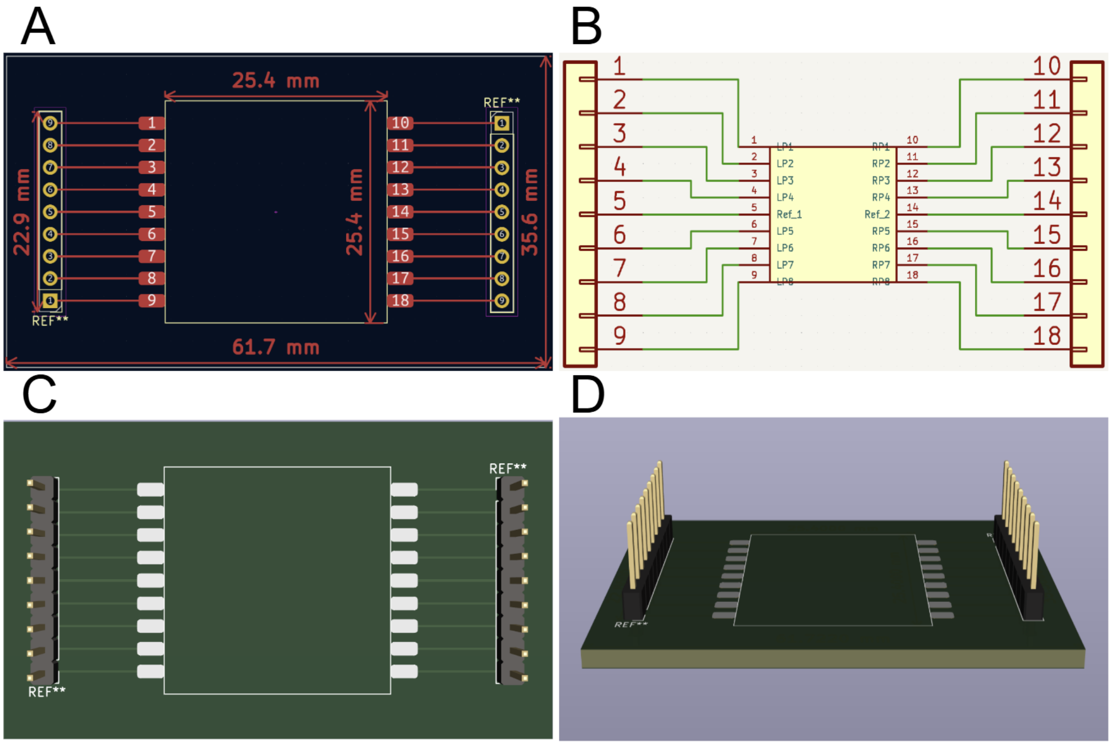


**Fig. S6. KiCAD design layout of printed circuit board (PCB) for the fully packaged shell MEAs. (A)** 2D representation of the PCB design of a 16-electrode PCB for the NO-integrated packaging. We positioned the reference electrodes on the edges of each electrode arrangement for ease of access. **(B)** 2D schematic representation showing the connectivity of the PCB with 18 bond pads. LP and RP refer to left and right pads, and Ref 1 and Ref 2 refer to reference electrodes. **(C)** The top view of the PCB: 61.7 mm in length and 35.6 mm in width. The square boundary that fits the MEA chip is 25.4 mm in length. **(D)** Side-angled view of the 3D representation of the PCB.

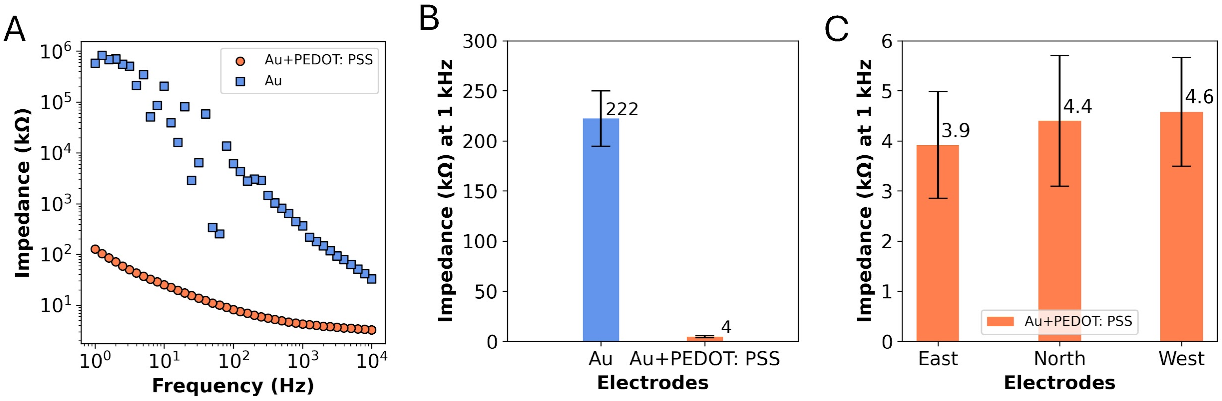


**Fig. S7. Electrochemical impedance measurements with PEDOT: PSS coated electrodes. (A)** Plot of the average (three samples) impedance as a function of frequency (1 Hz to 10 kHz) before and after PEDOT: PSS deposition. **(B)** At 1 kHz, the average impedance values are significantly lower when Au electrodes were coated with PEDOT: PSS. **(C)** Plot of the average impedance at 1 kHz for individual PEDOT: PSS-coated electrodes labeled East, West, and North, demonstrating consistent performance across all samples.

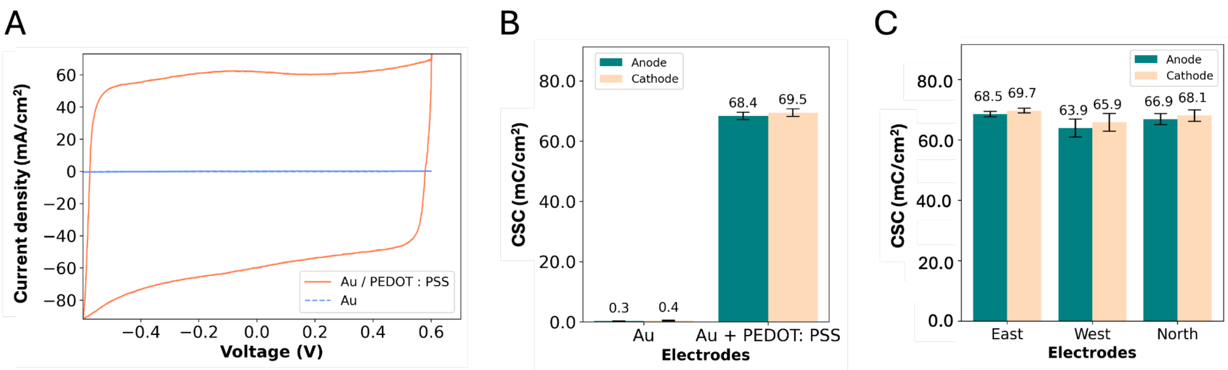


**Fig. S8. Charge storage capacity (CSC) measurements with PEDOT: PSS coated electrodes. (A)** Cyclic voltammetry using a 200 mV $s^{-1}$ scan rate depicting the anodal and cathodal CSCs of the Au electrode with and without PEDOT: PSS. **(B)** Plot of charge storage density at bare Au and PEDOT: PSS coated electrodes. The electrodeposited PEDOT: PSS electrode exhibited a statistically significant increase in CSC compared to the bare Au electrode. **(C)** Bar diagram showing the CSC characteristics of PEDOT: PSS coated East, West, and North electrodes illustrating consistent performance.

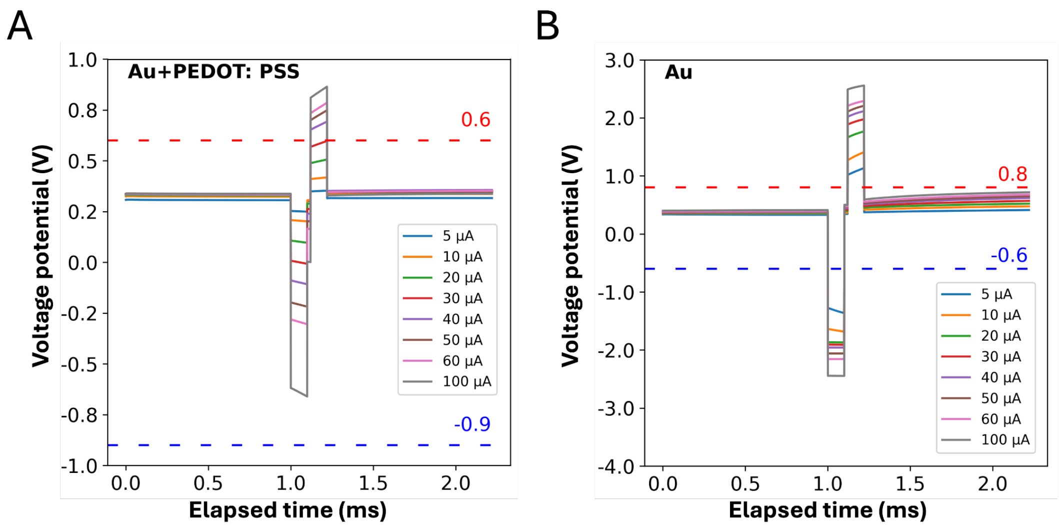


**Fig. S9. Voltage transients of contact pads.** Electrolysis windows for the electrodes are indicated by red and blue dotted lines for the anodal and cathodal limits, respectively (*1*). **(A)** PEDOT: PSS -coated Au electrodes can receive a significantly higher charge per unit area without the polarization potential crossing the water electrolysis window. **(B)** Bare Au electrode showed a higher cathodic and anodic polarization potential for the same range of current stimulation (5-100 µA), ultimately reaching the current injection capacity (CIC) at 100 µA.

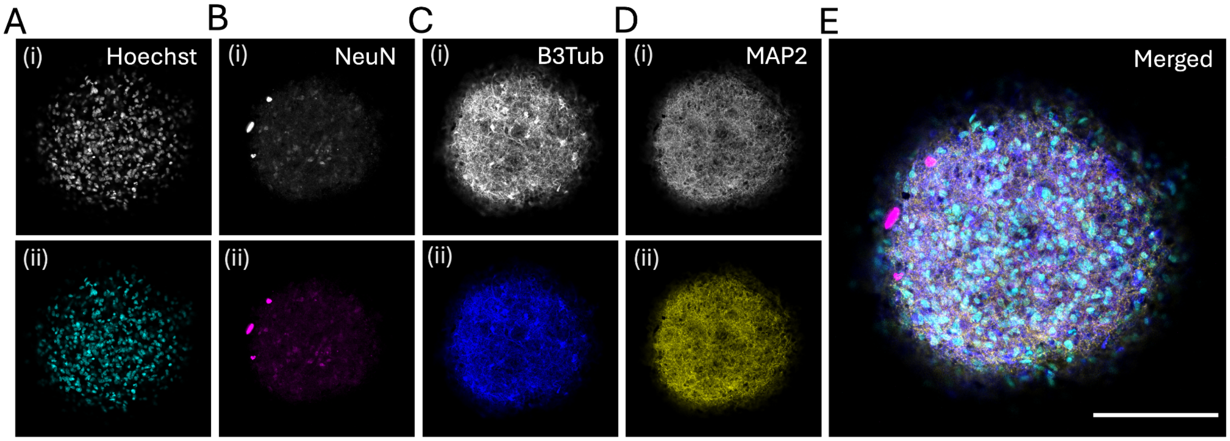


**Fig. S10. Expression of neuronal markers in eight-week-old organoids**. Confocal images of representative maximum intensity z-projections. **(A)** Presence of nuclei (Hoechst), **(B)** mature neurons (NeuN), **(C)** immature neurons (B3TUB), and **(D)** dendrites (MAP2) are shown in grayscale (i) and turquoise, purple, blue, and yellow, respectively (ii). **(E)** In the composite image, channels are overlaid to show colocalization of staining. Images were taken at 20x. Scale bar is 100 µm.

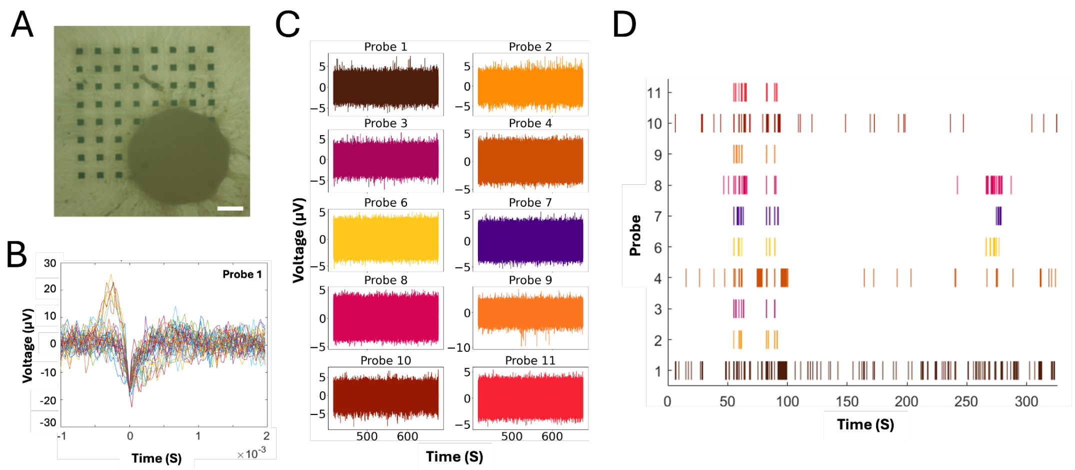


**Fig. S11. Raw recordings and extracted spike shape using commercial 2D MEAs. (A)** Brightfield microscopy image of 8-week differentiated NO on commercial MED 64 MEA plate (MED-P515A). Scale bar is 200 µm **(B)** Overlayed spike shape from a recording of an 8-week-old organoid extracted from probe 1 (a single electrode) on the 2D MEA shell, **(C)** Raw recordings of ten probes measured for five minutes. **(D)** Raster plot of corresponding ten probes filtered at five times the signal RMS voltage.

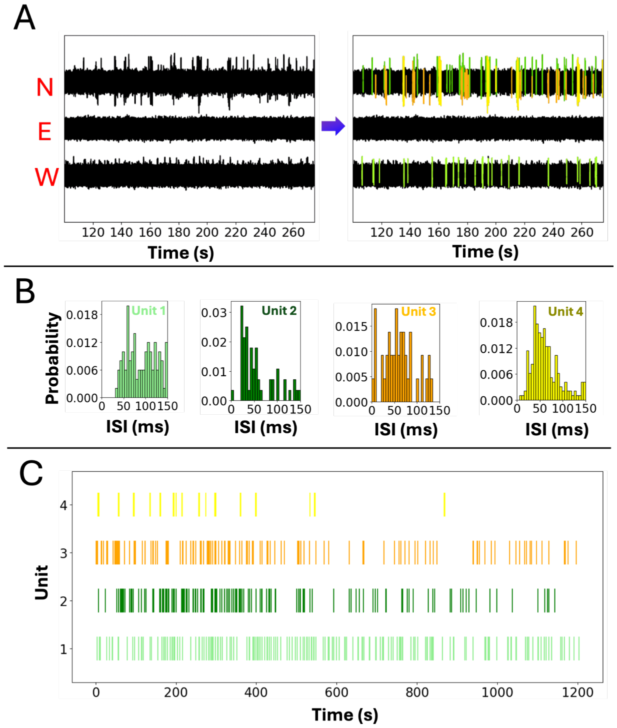


**Fig. S12. Spike time unit analysis. (A)** A snippet (120 and 260 s) of the recorded signals from organoid before and after highlighting the spikes generated by the North (N), East (E), and West (W) electrodes. Spikes are categorized as light green (Unit 1), orange (Unit 2), dark green (Unit 3), and yellow (Unit 4). **(B)** The probability distribution plot for interspike intervals (ISI) corresponds to Units 1, 2, 3, and 4. **(C)** Individual spike units, as shown in panel A, comparing spikes over time.

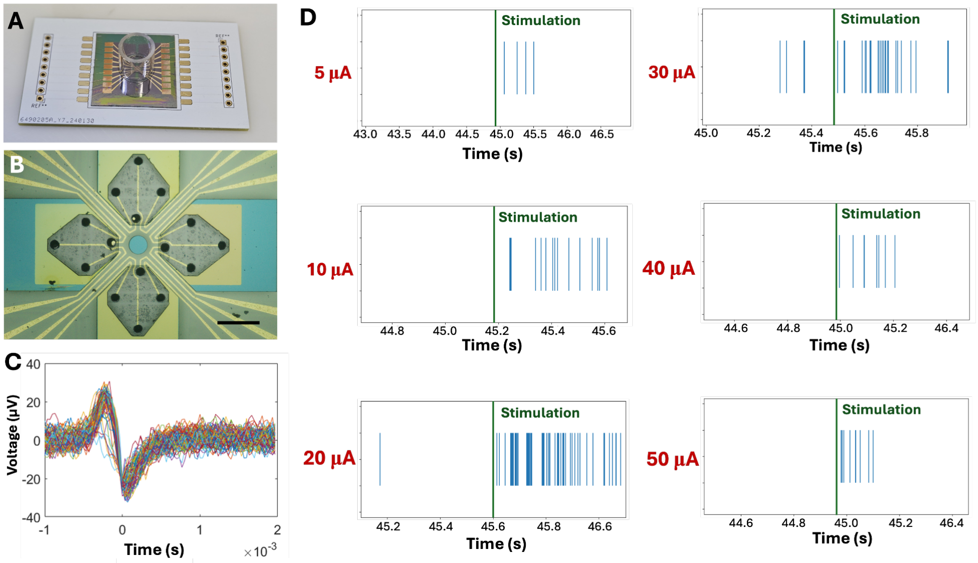


**Fig. S13. Integrated package with NO interfacing shell MEA and evoked response via electrical-induced stimuli. (A)** Image of 16-electrode shell MEA, PDMS bonded to a printed circuit board (PCB). **(B)** Brightfield microscopy of as-fabricated four-leaflet, 16-electrode, shell MEAs design with PEDOT: PSS coated electrodes. **(C)** Spike waveform plots of a typical single-unit cluster. **(D)** Organoid burst activity was observed at 0.5 s before and after stimulation at various current levels (5 µA to 50 µA). Stimulation currents of 20-30 µA showed the most frequent and dense spike activity, suggesting that this current elicits the greatest neuromodulation within the tested range.

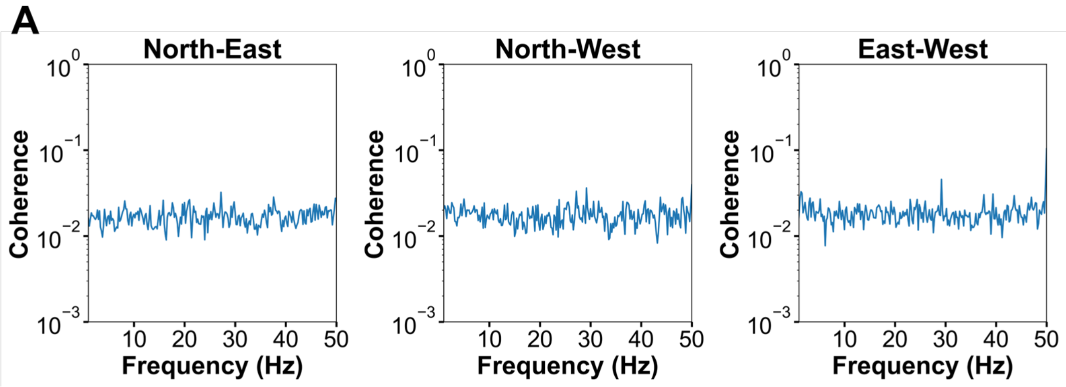


**Fig. S14. Pre-stimulation coherence analysis of NOs using three-electrode 3D shell MEAs. (A)** Coherence spectra of NO spontaneous activity calculated over a 200-second interval with a 5-second window size in the absence of stimulation. All electrode pairs exhibit consistently low coherence values (~0.02) across the 1-50 Hz frequency range, indicating weak synchronization and minimal functional connectivity between the examined regions.

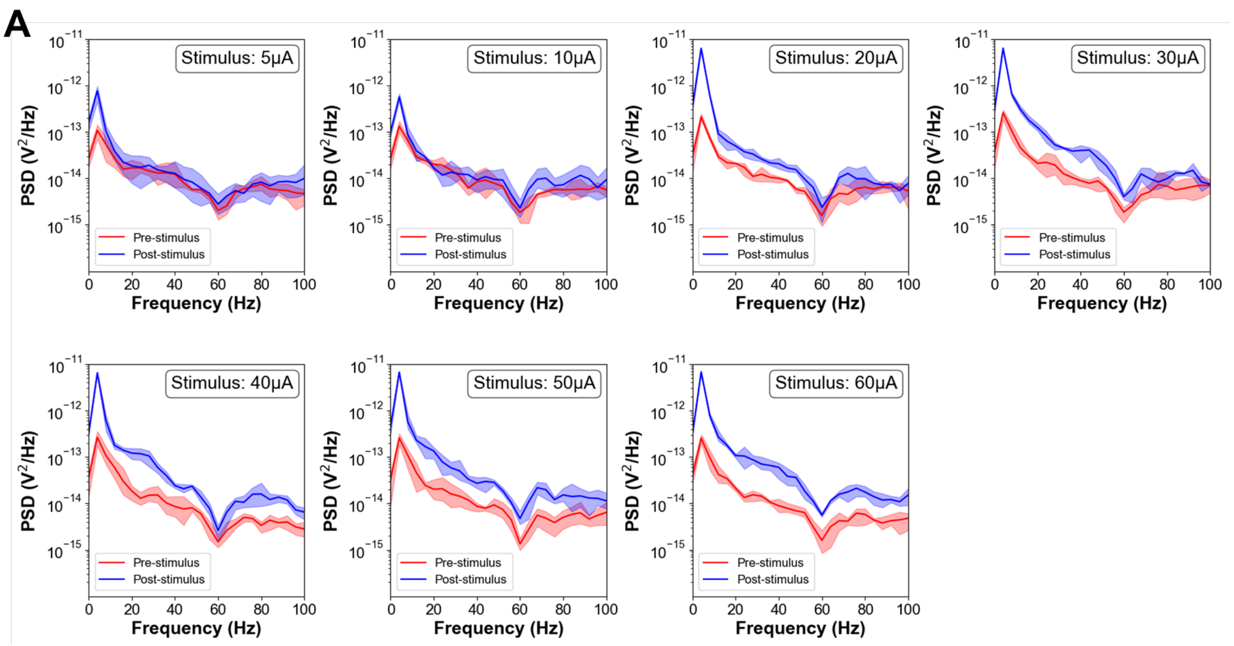


**Fig. S15. PSD analysis of NO responses to electrical stimulation at varying intensities. (A)** PSD spectra calculated for neural signals recorded pre- and post-stimulation at different current intensities (5 µA, 10 µA, 20 µA, 30 µA, 40 µA, 50 µA, and 60 µA). Each panel represents the average PSD across trials with shaded regions indicating the standard deviation. Post-stimulation signals (blue) exhibited a significant increase in power, particularly in the low-frequency range (1–10 Hz), compared to pre-stimulation signals (red). This enhancement in PSD intensity was more pronounced at all frequencies from 20 µA onwards. The results demonstrated the capacity of the shell MEAs to induce frequency-dependent changes in neural dynamics.

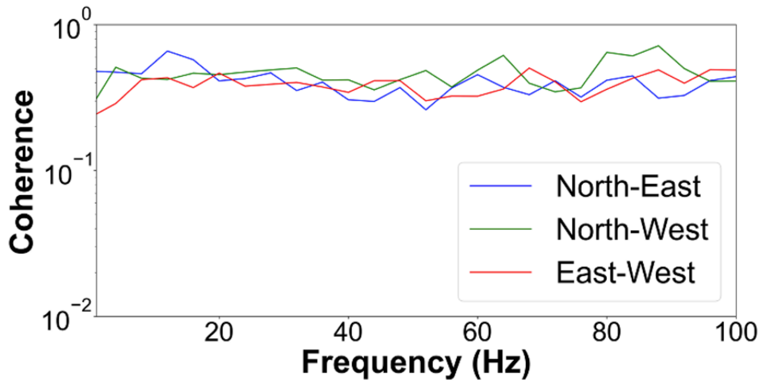


**Fig. S16. Coherence analysis measured using a silver bead (positive control) with high stimulating current using three-electrode shell MEA.** Coherence spectra for electrode pairs North-East, North-West, and East-West calculated from the stimulated signals of a silver bead within 1.5 seconds post-stimulation. Analysis was performed with a 0.25-second window size, using a stimulating current of 40 µA. The spectra confirmed high coherence (above 0.5) due to the shared phase reference.

#### Supplementary Note 1: Data Analysis

We collected all data output by the Intan RHS Stim/Recording System at a sampling rate of 30 kHz with either three or 16 active channels in the Intan-proprietary .rhs file format. The recordings were generally split based on day, organoid, and stimulation parameters, with an average recording time of about 10 minutes. We set a 60 Hz notch filter to compensate for electrical noise during recording.

We first chose organoids of interest to prepare this data for analysis based on the recorded activity. In Python-based Jupyter Notebooks, we extracted data from each of the .rhs files associated with the particular organoid(s) into a Pandas array. If the system did not record specific channels due to low data quality, we inserted zeros into the array as placeholders to ensure consistent data length and channel mapping for 16-channel data. Then, we saved the data and metadata, including electrode positions, stimulation waveforms, session information, start time/date, lab, and recording device, into the Neurodata Without Borders (NWB) (*2*) file format, allowing for compatibility with the SpikeInterface (*3*) Python library leveraged in the analysis.

Once ingested and formatted, we prepared the data for spike sorting by loading it into the SpikeInterface recording data structure and a custom ProbeInterface (*4*) model of either the three- or 16-channel shell. After, we set a bandpass filter with a 300-6000 Hz range and applied an optional whitening transformation (across the channels). Then, we performed a threshold-based artifact rejection at a 5-15 mV range. If a signal absolute value passed this threshold, we zeroed out in a window of 10 ms around the detected threshold.

With the data fully prepared, it was then input into the Mountainsort5 (*5*) spike sorting algorithm. First, events were detected using a default two-way threshold of 5.5 times the signal standard deviation. Based on the SNR of individual recorded signals and the metrics on individual unit clusters, the threshold was adjusted offline, creating a threshold ranging from 1.55 times to 11 times the signal standard deviation. For **Fig. 6**, the results of this detection were taken as MUA for visualization in 3D. We extracted features from the detected events using principal components analysis for all single-unit analyses, with three features per channel typically computed. We then clustered waveforms using the MountainSort5 approach, resulting in spike templates, a distribution of spike waveforms, and spike times for each detected cluster.

To verify the spike sorting results, we computed various visualizations and metrics. First, we plotted spike time rasters for all detected units against time. Next, we visualized each channel's 80 seconds of raw activity with a color-coded spike event—a plot of all assigned waveforms and the spike template for each cluster. Finally, we computed the inter-spike interval (ISI) distribution, including the minimum and maximum observed ISIs. Two researchers used these visualizations to ensure that clusters were not dominated by noise events or split into multiple events.

We reviewed clusters and detected events for spurious noise events or missed spikes evident in the raw trace. We re-evaluated the threshold, number of feature dimensions, and clustering in such cases—next, the analysis removed clusters dominated by noise events (lacking a consistent spike shape). Finally, if two clusters showed spike templates of the same shape and duration on the same electrode, it was identified as a split cluster. This cluster was merged into a single unit, incorporating spike waveforms and times. Both reviewers had to agree on removing noise clusters or merging clusters.

LFP analysis was conducted following a similar approach, using the NWB files for each recording. We used the SpikeInterface package to load each waveform. For artifact detection, we first band-pass filtered the signal between 300 and 6000 Hz and identified artifact times for single units. The raw signal was then bandpass filtered between 1 and 400 Hz, and the artifact times zeroed out of the signal and downsampled to 1000Hz. From here, we applied additional bandpass filters to investigate specific bands or used the Welch method (implemented in SciPy) directly to estimate power spectral densities. (*6*).

We conducted post-processing using the results of the spike sorting approach. If applicable, we loaded the NWB file for each waveform to extract the time of stimulation pulses. We used a window of 2 seconds before and after the stimulus to identify spike events for each unit, determine the spike rate before and after stimulation, calculate the difference in pre- and post-stimulation rates, and use it to compute histograms and box plots.

We used the spike sorting procedure results to analyze the firing behavior of detected units over time. For noise reduction, a moving average approach with a 10-second window was applied after initial exploration, balancing temporal resolution and noise reduction. In **Fig. 5C**, each star in the graph represents the number of firings for a detected unit: the first star corresponds to 0–10 seconds, the second to 1–11 seconds, and so on. Three stimulations occurred during the recording at 30, 45, and 60 seconds. Dashed lines in the graph indicate the adjusted locations of the stimulations. These adjustments account for the moving average window, which includes firing events up to 10 seconds into the future, thereby aligning the dashed lines with the proper recording times.

**Supplementary Note 2: Note on coherence analysis**

Coherence is a frequency-domain analysis tool that combines signal amplitude and phase information to measure the linear correlation of two signals at a particular frequency. For LFP signals, researchers can use coherence to study interactions between different organoid regions, network synchronization properties, and functional connectivity of neural activity. (*7*). High coherence (close to 1) indicates that two signals are strongly correlated, meaning that the signals change synchronously and may have similar amplitude patterns and phase relationships. Low coherence (close to 0) indicates that the two signals are not significantly correlated, meaning that the signals may be independent. We based the calculation of coherence on the cross-spectral densities $Sxy(f)$ and the respective power spectral densities $Sxx(f)$ and $Syy(f)$ of the two signals $x(t)$ and $y(t)$. The formula is

$$Cxy(f) = | Sxy(f) |^2 / (Sxx(f)\, Syy(f)).$$

Since the calculation process depends on the frequency domain characteristics of the signals, it is necessary first to estimate the spectral density using the Welch method (*6*). In this experiment, the signal preprocessing method for coherence analysis was the same as that for LFP analysis**,** and we calculated coherence via the coherence function of the SciPy.signal module in Python. The coherence function uses the Welch method, which divides the signal into multiple segments with 50% overlap, applies a window function to each segment to reduce spectral leakage, calculates the PSD of each signal and the cross power spectral density (CPSD) between them, and then averages these estimates across segments to obtain a smoothed and reliable coherence estimate. The coherence plots before and after stimulation for each electrode pair (N-E, N-W, and E-W) were the average results calculated from 20 trials.

#### Supplementary Note 3: Note on spatial stimulation and recording.

We detected multi-unit spike events by first band-pass filtering each channel between 300 and 6000Hz. Then, we detected events using the same procedure as the spike sorting with SpikeInterface. We analyzed windowed pre/post task spike sorted firing counts utilizing a window and counting spike events for each unit identified from spike sorting within this window. We used fixed pre- and post-stimulus windows (with a buffer to account for stimulation artifacts) and a non-overlapping rolling time window. In addition to spike times, we utilized a similar approach for channel-level statistics to estimate the spatial effects of stimulation. Given a presumed organoid radius, we fold each electrode in idealized unit spherical arcs around each organoid. Letting $E_k$ being the statistic value (such as spike count, LFP, etc.) at electrode $k$, we will consider the smoothed value at position $P$ as the simplicial average of

$$w(E_k, P) = \phi\{d(E_k, P/\sigma\}/(\Sigma\{d(E_k, P/\sigma\}),$$

which is the normalized weight and $\phi$ is a standard Gaussian kernel. Distance is then calculated as $d(E_k, P) = ||S_k{'}S||$where $S_{k'}$ is the position of $E_{k'}$ on the unit sphere and $S$ is the associated position of $P$. **Fig. 6** in the main text represents the smoothing where the value is the log of the post-stimulus exceedance counts in a 5-second window with the 16-electrode shell system.

**Supplementary Table 1.** Primary and secondary antibodies used for immunohistochemistry.

| **Antibody** | **Host** | **Type** | **Source** | **Catalog number** | **Dilution** |
| --- | --- | --- | --- | --- | --- |
| NeuN | Mouse | Monoclonal | Phospho | 583-FOX3 | 1:500 |
| B-III-Tubulin | Rabbit | Monoclonal | Cell signaling | 5568 | 1:1500 |
| MAP2 | Chicken | Polyclonal | Invitrogen | PA1-10005 | 1:5000 |
| SOX2 | Rat | Monoclonal | Invitrogen | 14-9811-82 | 1:100 |
| NF-H | Chicken | Polyclonal | Invitrogen | PA1-10002 | 1:5000 |
| Alexa Fluor$^{TM}$ 488 goat anti-rat IgG (H+L) | Rat | Polyclonal | Invitrogen | A-48262 | 1:200 |
| Alexa Fluor$^{TM}$ 488 goat anti-rabbit IgG (H+L) | Rabbit | Polyclonal | Invitrogen | 35552 | 1:500 |
| Alexa Fluor$^{TM}$ 568 goat anti-mouse IgG (H+L) | Goat | Polyclonal | Invitrogen | A11001 | 1:500 |
| Alexa Fluor$^{TM}$ Plus 647 goat anti-chicken IgY (H+L) | Goat | Polyclonal | Invitrogen | A32933 | 1:500 |
